# Structure-based design of a soluble human cytomegalovirus glycoprotein B antigen stabilized in a prefusion-like conformation

**DOI:** 10.1101/2024.02.10.579772

**Authors:** Madeline R. Sponholtz, Patrick O. Byrne, Alison G. Lee, Ajit R. Ramamohan, Jory A. Goldsmith, Ryan S. McCool, Ling Zhou, Nicole V. Johnson, Ching-Lin Hsieh, Megan Connors, Krithika Karthigeyan, John D. Campbell, Sallie R. Permar, Jennifer A. Maynard, Dong Yu, Matthew J. Bottomley, Jason S. McLellan

## Abstract

Human cytomegalovirus (HCMV) glycoprotein B (gB) is a class III membrane fusion protein required for viral entry. HCMV vaccine candidates containing gB have demonstrated moderate clinical efficacy, but no HCMV vaccine has been FDA-approved. Here, we used structure-based design to identify and characterize amino acid substitutions that stabilize gB in its metastable prefusion conformation. One variant containing two engineered interprotomer disulfide bonds and two cavity-filling substitutions (gB-C7), displayed increased expression and thermostability. A 2.8 Å resolution cryo-electron microscopy structure shows that gB-C7 adopts a prefusion-like conformation, revealing additional structural elements at the membrane-distal apex. Unlike previous observations for several class I viral fusion proteins, mice immunized with postfusion or prefusion-stabilized forms of soluble gB protein displayed similar neutralizing antibody titers, here specifically against an HCMV laboratory strain on fibroblasts. Collectively, these results identify initial strategies to stabilize class III viral fusion proteins and provide tools to probe gB-directed antibody responses.

**Teaser:** A structure-based design campaign leads to stabilization of the class III viral fusion protein from HCMV in a prefusion-like conformation.

## Introduction

Human cytomegalovirus (HCMV), also known as human betaherpesvirus 5 (*1*), establishes lifelong latency in infected individuals (*2*). HCMV infects 60–90% of adults worldwide and can be spread through transplacental transmission from mother to fetus or through contact with bodily fluids (*2, 3*). Primary infection, reinfection, and reactivation of a latent infection all pose risks for the fetus during pregnancy (*2*). HCMV is the leading infectious cause of birth defects worldwide (*4*), with about 1 in 200 newborns having congenital infections in the US (*5*). HCMV is also a common and serious opportunistic infection following solid-organ and stem-cell transplants (*2*). In healthy patients, HCMV infection can be controlled by a robust immune response, which may come at the cost of decreased immune function over time (*6*). Despite the considerable disease burden associated with HCMV and the variety of vaccine formulations investigated over the last five decades, no FDA-approved vaccine is available for prevention or treatment (*7*).

As a member of the *Herpesviridae* family, HCMV is an enveloped, double-stranded DNA virus (*8*). Viruses within this family enter host cells through a conserved mechanism, relying on the coordinated actions of multiple surface glycoproteins for receptor binding and membrane fusion (*9*). The specific receptors and glycoproteins involved vary among different herpesviruses, although glycoprotein B (gB), which mediates membrane fusion, is highly conserved and required for entry (*10, 11*). For HCMV, receptor binding is mediated by glycoprotein complexes referred to as Trimer (gH, gL, and gO) and Pentamer (gH, gL, UL128, UL130, and UL131A) (*12–14*). Receptor binding to Trimer and Pentamer complexes is thought to trigger the irreversible transition of gB from a metastable prefusion conformation to a highly stable postfusion conformation, thereby facilitating fusion of the viral and host-cell membranes. Given its essential role in viral entry, gB is typically included in HCMV vaccine candidate formulations (*15*). Of the vaccine candidates tested in humans, recombinant gB delivered with the MF59 adjuvant has shown promise in phase II clinical trials (NCT00125502, NCT00133497), achieving 40– 50% short-lived efficacy in preventing HCMV infection in both adolescent and postpartum cohorts (*16, 17*), as well as reducing viremia and antiviral prophylaxis in renal transplant patients (*18*). The immune correlate of protection against HCMV acquisition from these studies was plasma immunoglobulin G (IgG) binding to native gB expressed on the surface of a cell (*19*), suggesting that the conformation of gB is critical to vaccine efficacy.

HCMV gB is a class III fusion protein encoded by open reading frame UL55. Class III fusion proteins share features of both class I and class II fusion proteins, including trimeric central helices arranged in a coiled-coil and internal fusion loops (*9, 20*). Class III fusion proteins are found in herpesviruses, rhabdoviruses, thogotoviruses, and baculoviruses (*9*). Upon translation of the monocistronic gB mRNA, each protomer undergoes extensive glycosylation, with high occupancy at the 17–18 predicted N-linked glycosylation sites per protomer (*21*) (**Figure 1A**). Additionally, two O-linked glycosylation sites are located near the N-terminus of each protomer (*22*). Three gB protomers associate to form a metastable prefusion trimer, which is processed by furin (457-R-X-K/R-R-460) and incorporated into the HCMV virion. Each protomer has two hydrophobic fusion loops located at the membrane-proximal tip of structural domain I (DI) that pack against the hydrophobic membrane-proximal region (MPR) of a neighboring protomer in the prefusion trimer (**Figures 1B**, **S1**) (*23*). The metastable prefusion gB trimer undergoes significant conformational rearrangement to facilitate membrane fusion. This involves transitioning to an extended intermediate state, in which DI disengages from the MPR, rotates with DII approximately 180° relative to DIII, and embeds the fusion loops into the host-cell membrane (*23*). This intermediate, which bridges the viral and host-cell membranes, then collapses into the highly stable postfusion conformation, bringing the viral and host-cell membranes together to create a fusion pore (*9, 23*).

**Figure 1.**
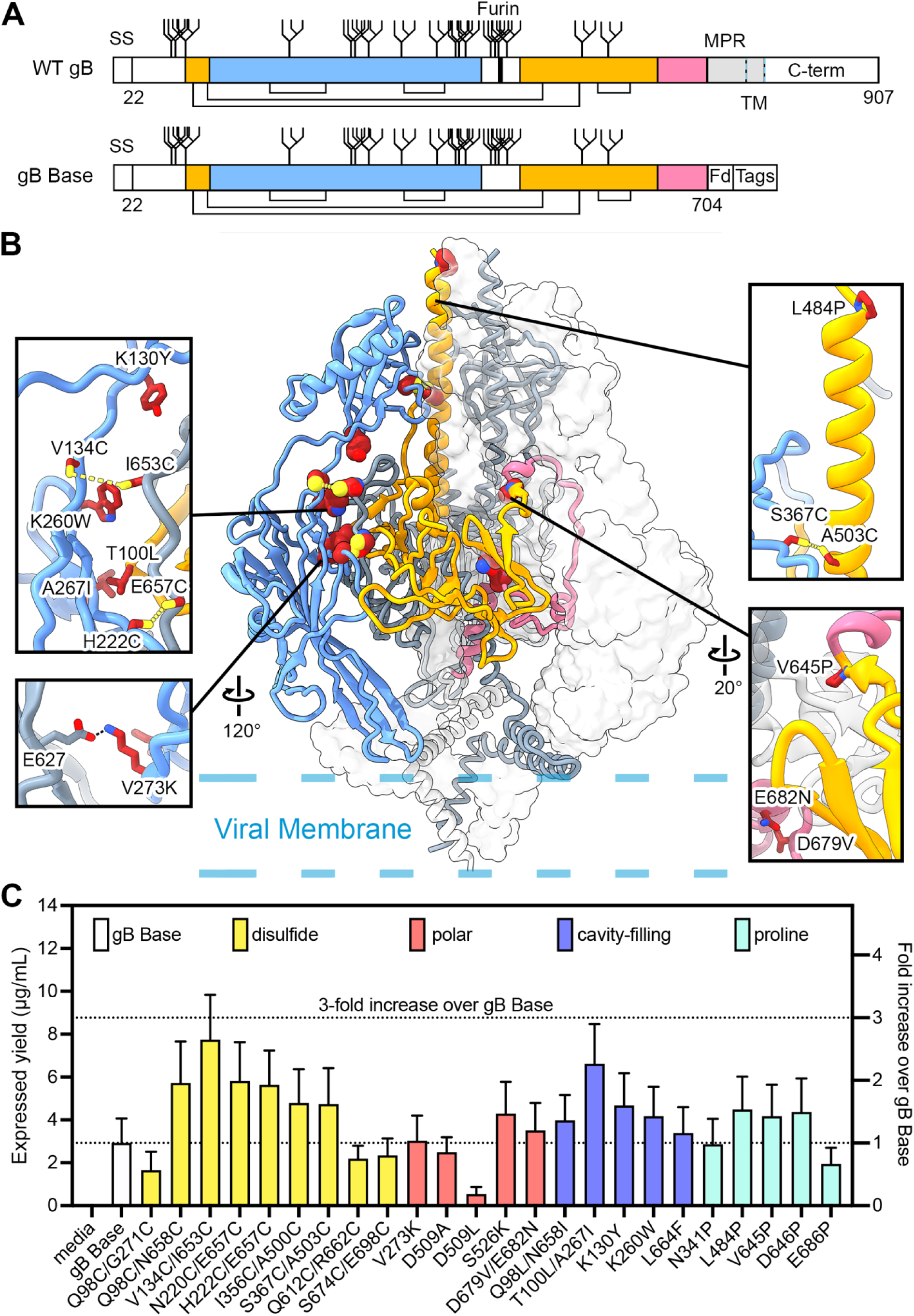
Exemplary substitutions for HCMV gB stabilization. (**A**) Schematic of wild-type (WT) Towne strain HCMV gB and ectodomain base construct (gB Base). Native disulfide bonds are shown as connecting black lines. N-linked glycosylation sites are shown as branched lines. The native furin cleavage site is shown as a thick black line. The N-terminal signal sequence (SS) and C-terminal domain (C-term) are shown as white boxes. The first and second regions expected to move during the conformational rearrangement from pre-to-postfusion gB are colored blue and pink, respectively, and the region that does not undergo substantial rearrangement is colored yellow. The membrane-proximal region (MPR) and transmembrane domain (TM) are shown in gray, with the TM demarcated by dashed blue and black lines. The soluble gB Base construct consists of the first 704 residues of WT gB with substitutions C246S, R457S, and R460S, of which the latter two remove the native furin cleavage site. In gB Base, residue 704 is followed by the foldon (Fd) domain and C-terminal tags. (**B**) Side view of trimeric prefusion HCMV gB (PDB ID: 7KDP) (*23*). One protomer is colored as in A and shown as a ribbon diagram, one protomer is colored gray and shown as a cartoon trace of the α-carbon backbone, and one protomer is shown as a transparent surface. Exemplary substitutions are shown as spheres with sulfur atoms in yellow, nitrogen atoms in blue, and oxygen atoms in red. The approximate location of the viral membrane is shown as dashed blue lines. Insets show exemplary substitutions as sticks. Anticipated hydrogen bonds are shown as black dashed lines and anticipated disulfide bonds are shown as yellow dashed lines. (**C**) Absolute and relative expression levels of individual variants determined by quantitative bio-layer interferometry (BLI). Variants are colored by substitution type.

Two 3.6 Å resolution crystal structures of postfusion HCMV gB were published in 2015 (*24, 25*) and these closely resembled the postfusion structures of gB homologs from herpes simplex virus type 1 (HSV-1) and Epstein-Barr virus (EBV) (*26, 27*). Subsequent cryo-electron tomography (cryo-ET) reconstructions of full-length, membrane-bound prefusion HCMV gB (*28–30*) revealed a shorter and more compact structure than observed for postfusion gB and suggested a domain architecture similar to prefusion G from vesicular stomatitis virus (VSV) (*31*). Recently, a 3.6 Å resolution cryo-electron microscopy (cryo-EM) structure of prefusion HCMV gB was determined using detergent-solubilized full-length gB purified from virions, complexed with the neutralizing human antibody SM5-1 (*32*), and stabilized in the prefusion conformation with a thiourea fusion inhibitor and a chemical cross-linker (*23*). This was the first high-resolution structure of any prefusion herpesvirus gB protein, providing the necessary data to guide the structure-based design of a stabilized prefusion immunogen.

In evaluating the humoral immune response to HCMV gB, antigenic mapping has identified six antigenic domains (AD-1–6), of which all but AD-3 and AD-6 are capable of eliciting neutralizing antibodies (*32, 33*). AD-1, located on structural domain IV (DIV) (**Figure S1**), is considered the immunodominant region of gB and elicits primarily non-neutralizing antibodies (*32, 34*). AD-1 is partially obscured by DI, DII and DV in the prefusion conformation but is highly accessible in the postfusion conformation (*23–25*). AD-2 consists of the first 85 N-terminal residues of gB, which are flexible and have not been resolved in any HCMV gB structure but can elicit potently neutralizing antibodies that bind linear epitopes, including the human antibodies TRL345 and 3-25 (*35, 36*). AD-3 corresponds to the C-terminal cytoplasmic tail that is presumably inaccessible on intact virions and elicits exclusively non-neutralizing antibodies (*32, 37*). AD-4, located on DII, contains the epitope for broadly neutralizing antibodies 7H3 and SM5-1 (*32, 38*). AD-5, located on DI, contains the epitope for the neutralizing antibody 1G2 (*32*) and can elicit comparatively high titers of neutralizing antibodies (*39*). An additional antigenic region corresponding to DV, AD-6, was recently identified in a study assessing the antibody response elicited by the recombinant gB vaccine tested in clinical trial NCT00299260 and shown to elicit antibodies that are non-neutralizing but limit cell-to-cell spread of HCMV in both fibroblasts and epithelial cells (*33*). Although it has been shown that some prefusion-stabilized class I fusion proteins elicit higher quality immune responses relative to immunization with postfusion or non-prefusion-stabilized variants (*40–42*), this has not yet been reported for class III herpesvirus fusion proteins, primarily due to the difficulty of producing prefusion gB.

Here, we used the published structure of full-length, detergent-solubilized prefusion HCMV gB (PDB ID: 7KDP) (*23*) to guide the engineering of a soluble HCMV gB ectodomain construct stabilized in a prefusion-like conformation. We designed and biochemically characterized a soluble postfusion ectodomain base construct (gB Base) and numerous amino acid substitution variants. As no antibodies that bind exclusively to the prefusion conformation of HCMV gB have been described, we analyzed protein expression, thermal stability, monodispersity, and conformational homogeneity to evaluate gB variants. This led us to identify gB-C7, a variant with improved expression and thermostability relative to gB Base. A 2.8 Å resolution cryo-EM structure of gB-C7 in complex with neutralizing antibodies 1G2 and 7H3 (*32, 38*) reveals that gB-C7 folds into a prefusion-like conformation, even though it lacks the MPR and transmembrane domain. Mice were immunized with gB-C7 or the gB Base postfusion construct to assess the resulting gB-specific and HCMV-neutralizing antibody titers in fibroblasts. This work identifies a strategy for stabilizing class III viral fusion proteins, provides structural insights into the prefusion conformation of HCMV gB when bound by neutralizing antibodies, and creates new reagents for the isolation of gB-directed antibodies and assessment of antibody responses from infected or vaccinated individuals.

## Results

### Design and initial characterization of single substitution HCMV gB variants

We first designed a soluble base construct (gB Base) comprising the HCMV gB ectodomain (Towne strain, residues 1–704) followed by a C-terminal T4 fibritin (foldon) trimerization motif, an octa-histidine tag, and a Twin-Strep affinity tag (**Figure 1A**, **Figure S1**). We also substituted an unpaired cysteine at position 246 for serine (C246S) and eliminated the furin cleavage site with two serine substitutions (R457S, R460S), as described previously (*24*). Approximately 3 mg/L of gB Base was routinely purified from transiently transfected FreeStyle 293 cell cultures. To structurally characterize the base construct, we determined a 3.4 Å resolution cryo-EM structure of gB Base in complex with the fragments of antigen binding (Fabs) from high affinity, neutralizing antibodies 1G2 and 7H3 (*32, 38*) (**Figure S2**, **Table S1**). As expected, the structure revealed that gB Base adopts the postfusion conformation and closely resembles previously determined crystal and cryo-EM structures of postfusion HCMV gB (**Figure S3**) (*23–25*).

To stabilize the soluble gB ectodomain in the prefusion conformation, we designed and characterized 24 gB variants, each containing one or two amino acid substitutions (**Figure 1B**). Types of substitutions included: engineered disulfide bonds to covalently link regions that separate during the conformational rearrangement from prefusion to postfusion (disulfide), substitutions to neutralize internal charge imbalances (polar), hydrophobic residues to fill internal cavities (cavity-filling), and proline residues to disfavor refolding of secondary structure (proline) (*43*). To evaluate the expression of the variants, we performed small-scale (4 mL) transient transfections of FreeStyle 293 cells followed by quantification of the expressed protein yield in clarified media by bio-layer interferometry (BLI, **Figure 1C**). The gB Base construct yielded 2.9 μg/mL of medium on average and the variants yielded between 0.5 and 7.7 μg/mL. Of the 24 variants tested, 16 increased HCMV gB expression relative to gB Base (six disulfide, two polar, five cavity-filling, and three proline).

We next purified variants from 40 mL cultures of FreeStyle 293 cells to conduct further characterization. When purified and analyzed by reducing SDS-PAGE, gB Base and its variants yielded a predominant band around 130 kDa (**Figures 2A**, **S4**), consistent with the molecular weight of glycosylated monomeric HCMV gB ectodomain. When analyzed by non-reducing SDS-PAGE, gB samples migrated as three distinct species, likely corresponding to monomers and disulfide-linked dimers and trimers (**Figure 2A**). Five interprotomer disulfide variants (Q98C/N658C, V134C/I653C, N220C/E657C, H222C/E657C, and S674C/E698C) exhibited the greatest proportion of the trimer species (bands >460 kDa), indicating the formation of intermolecular disulfide bonds between protomers. As expected, disulfide variants designed to form intramolecular bonds predominately ran as monomers (**Figure 2A**).

**Figure 2.**
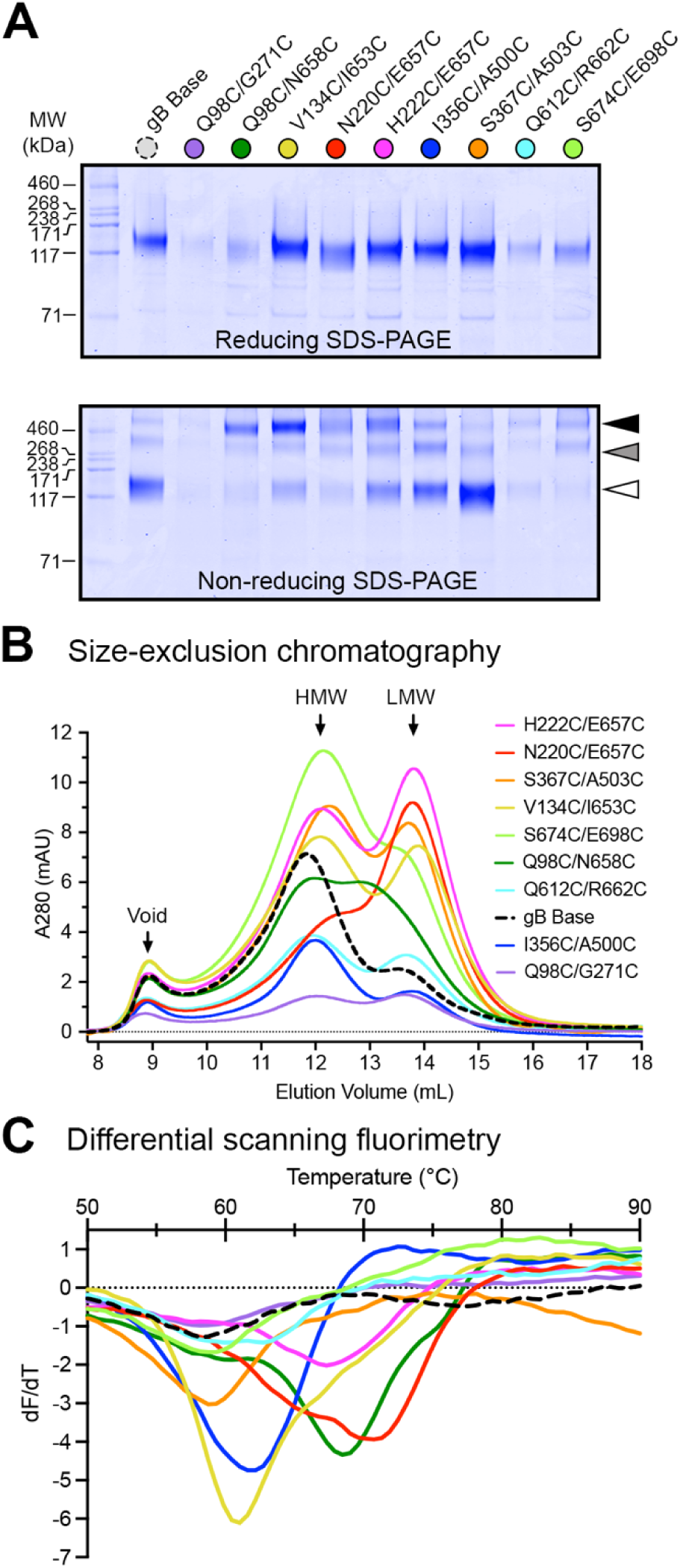
Characterization of single disulfide gB variants. (**A**) Reducing and non-reducing SDS-PAGE of gB variants. Molecular weight standards are indicated at the left in kDa. White, gray, and black triangles correspond to monomeric, dimeric, and trimeric gB molecular weights, respectively. (**B**) Size-exclusion chromatography (SEC) traces of purified gB variants. The approximate location of the void volume, high molecular weight (HMW) peak, and low molecular weight (LMW) peak are identified with black arrows. (**C**) Differential scanning fluorimetry (DSF) analysis of gB variant thermostability colored as in B.

All single disulfide variants were then analyzed by size-exclusion chromatography (SEC) and differential scanning fluorimetry (DSF) to assess their solution behavior and thermal stability. When separated by SEC, the gB Base construct yielded two main peaks: one high molecular weight (HMW) peak with a retention volume of ∼12 mL and one low molecular weight (LMW) peak with a retention volume of ∼14 mL (**Figure 2B**). The LMW peak was consistent with gB ectodomain trimers, whereas the HMW peak was consistent with oligomers of trimers, which are known to form via the exposed hydrophobic residues in the fusion loops (*25, 44*). Since the hydrophobic fusion loops are farther apart in the prefusion conformation of gB (*23*), we hypothesized that stabilizing gB in its prefusion conformation would disrupt the hydrophobic oligomer interface. This, in turn, would disfavor higher-order oligomerization and favor monodisperse trimers, quantified as an increase in the ratio of the area under the curve (AUC) of the LMW peak relative to the HMW peak (AUC_LMW/HMW_). The gB Base construct displayed a low AUC_LMW/HMW_ ratio, indicating fewer monodisperse trimers, whereas a subset of interprotomer disulfide variants exhibited relatively high AUC_LMW_/_HMW_ ratios, indicating more monodisperse trimers (**Table S2**). As evaluated by DSF, five disulfide variants displayed increases in melting temperature (T_m_) compared to the base construct (T_m_ of 59 °C): variants V134C/I653C, I356C/A500C, H222C/E657C, Q98C/N568C, and N220C/E657C exhibited T_m_ values of 61, 62, 67, 69, and 71 °C, respectively (**Figure 2C**). Of these, all but I356C/A500C were interprotomer disulfide variants. Overall, interprotomer disulfide variants V134C/I653C, N220C/E657C, and H222C/E657C displayed pronounced increases in expressed protein yield (**Figure 1C**), monodispersity (**Figure 2B**), and thermal stability (**Figure 2C**) relative to gB Base.

### Biochemical and structural characterization of combination variants

We next engineered seven gB combination variants containing two or three beneficial single substitutions and evaluated them for additive effects (listed in **Table 1**). We first selected V134C/I653C, N220C/E657C, and H222C/E657C based on their increased expression, favorable AUC_LMW_/_HMW_ ratios, and T_m_ values (**Figures 1C**, **2B**, **C**). Notably, all three disulfides were designed to covalently link DI and DV, which are adjacent in the prefusion conformation but separated in the postfusion conformation (**Figures 1B**, **S1**, **S3**). We selected four additional variants (T100L/A267I, K130Y, K260W, and V273F) based on their increased expression relative to gB Base (**Figure 1C**). As judged by quantitative BLI, all seven gB combination variants (gB-C1 through gB-C7) increased expression relative to gB Base (**Figure 3A**). Variants gB-C1 (N220C/E657C, T100L/A267I), gB-C6 (H222C/E657C, V134C/I653C), and gB-C7 (H222C/E657C, V134C/I653C, T100L/A267I) displayed additive increases in expression relative to gB Base and their constituent single substitution variants (**Figures 1C**, **3A**). In contrast, gB-C4 (N220C/E657C, K130Y) exhibited slightly reduced expression relative to the N220C/E657C variant, suggesting these substitutions may interfere with each other (**Figures 1C**, **3A**). When analyzed by non-reducing SDS-PAGE, all combination variants displayed an increased proportion of the trimer species relative to gB Base (**Figure 3B**). DSF analysis revealed that five combination variants exhibited increases in thermal stability relative to their parental single disulfide variants; gB-C6, gB-C5, and gB-C7 exhibited T_m_ values of 71, 72, and 72 °C, respectively (**Figure 3C**). In addition, gB-C4 and gB-C1 exhibited biphasic DSF profiles with dominant peaks at 73 and 75 °C, respectively, and shoulders around 68 °C. Biphasic DSF profiles may reflect a heterogeneous population with two unique unfolding events (*45*), or a two state transition, such as the dissociation of a higher order oligomer followed by the denaturation of the trimeric fusion protein (*46, 47*), neither of which were considered favorable.

**Figure 3.**
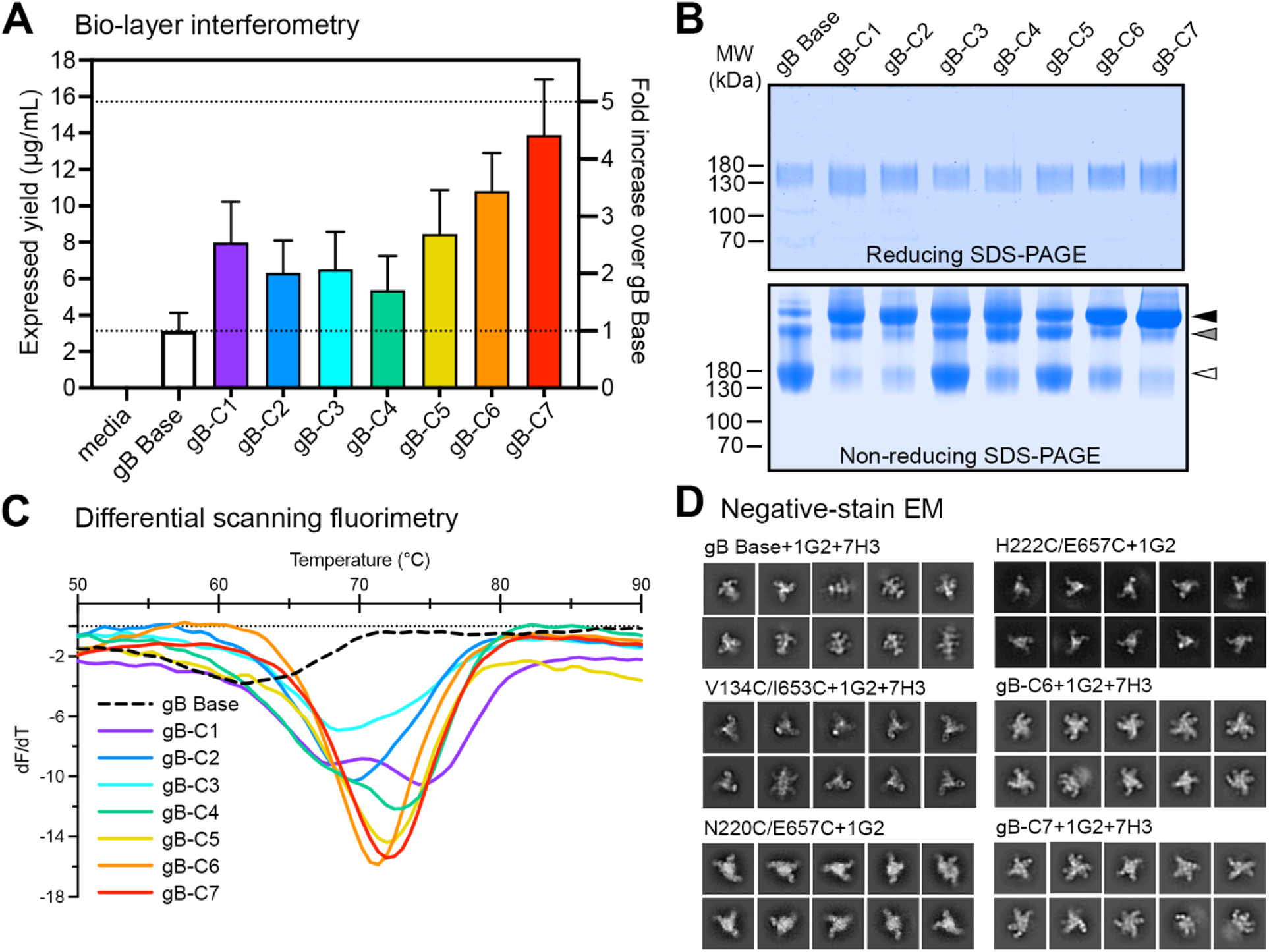
Characterization of combination gB variants. (**A**) Absolute and relative expression levels of individual variants determined by quantitative bio-layer interferometry (BLI). Dashed lines denote 1-fold and 5-fold increases in expression relative to gB Base. (**B**) Reducing and non-reducing SDS-PAGE gels of gB variants. Molecular weight standards are indicated at the left in kDa. White, gray, and black triangles correspond to monomeric, dimeric, and trimeric gB molecular weights, respectively. (**C**) Differential scanning fluorimetry (DSF) analysis of gB variant thermostability colored as in A. (**D**) Negative-stain EM 2D class averages of six variants.

**Table 1.** HCMV gB combination variants.

| **Combo variant name** | **Substitution 1** | **Substitution 2** | **Substitution 3** |
| --- | --- | --- | --- |
| gB-C1 | N220C/E657C | T100L/A267I | - |
| gB-C2 | H222C/E657C | K260W | - |
| gB-C3 | H222C/E657C | V273F | - |
| gB-C4 | N220C/E657C | K130Y | - |
| gB-C5 | N220C/E657C | V134C/I653C | - |
| gB-C6 | H222C/E657C | V134C/I653C | - |
| gB-C7 | H222C/E657C | V134C/I653C | T100L/A267I |

To assess the effect of single and multiple substitutions on the conformation of the soluble HCMV gB ectodomain, we conducted structural studies on a subset of variants by negative-stain EM (ns-EM) (**Figures 3D**, **S5**). We used two high-affinity, neutralizing antibodies to aid in conformational characterization, 1G2 and 7H3, which bind DI and DII, respectively (*32, 38*). Complexes of the gB Base construct with 1G2 and 7H3 yielded 2D class averages consistent with gB in the postfusion conformation bound to three 1G2 Fabs and either zero, one, or two 7H3 Fabs (**Figures 3D**, **S5**). Oligomers of postfusion gB (also known as rosettes) were observed for gB Base by ns-EM, consistent with the HMW peaks observed by SEC and previous findings on the spontaneous oligomerization of soluble postfusion gB (**Figure S5**) (*25, 44, 48*). The single disulfide variants V134C/I653C, N220C/E657C, and N222C/E657C each appeared more compact than gB Base, although a minority of the 2D class averages corresponded with side views of gB in the postfusion conformation (**Figures 3D**, **S5**). We hypothesized that these more compact particles corresponded to gB in prefusion-like conformations. Although single substitution variants T100L/A267I and S367C/A503C appeared promising based on their increased expression relative to gB Base (**Figure 1C**), both appeared to be in the postfusion conformation by ns-EM (**Figure S5**). The combination variants gB-C6 and gB-C7, both of which contain the V134C/I653C substitution, appeared to be conformationally similar to the single disulfide V134C/I653C variant (**Figure 3D**). Additionally, gB-C7 displayed distinct side views compared to gB Base (**Figure 3D**). The gB-C7 variant contains two disulfide substitutions along with a paired cavity-filling substitution (V134C/I653C, H222C/E657C, T100L/A267I). Overall, gB-C7 displayed higher expression than gB Base by a factor of 4.2, exhibited the greatest trimeric fraction when analyzed by non-reducing SDS-PAGE, had an 11 °C increase in T_m_ relative to gB Base, and resembled prefusion gB by ns-EM (**Figures 3A**–**D**). Given these properties, we focused on this construct for additional characterization.

### HCMV gB-C7 maintains a prefusion-like conformation

We determined a cryo-EM structure of gB-C7 in complex with 1G2 and 7H3 Fabs that reached a global resolution of 2.8 Å when refined with C3 symmetry (**Figures 4**, **S6**, **S7**, **Table S1**). We performed local refinement to account for relative motion between DI bound by 1G2 and the rest of the gB-C7 complex, yielding a 3.1 Å resolution local reconstruction (**Figure S6**). We then combined the 2.8 Å global map with the 3.1 Å local map in Phenix to generate a composite map (*49*), which we used to model the gB-C7 complex (**Figure 4A**). With this high-resolution map, we were able to model previously unresolved residues at the gB membrane-distal apex (residues 437–447 and 474–482, **Figures 4C**, **S8**).

**Figure 4.**
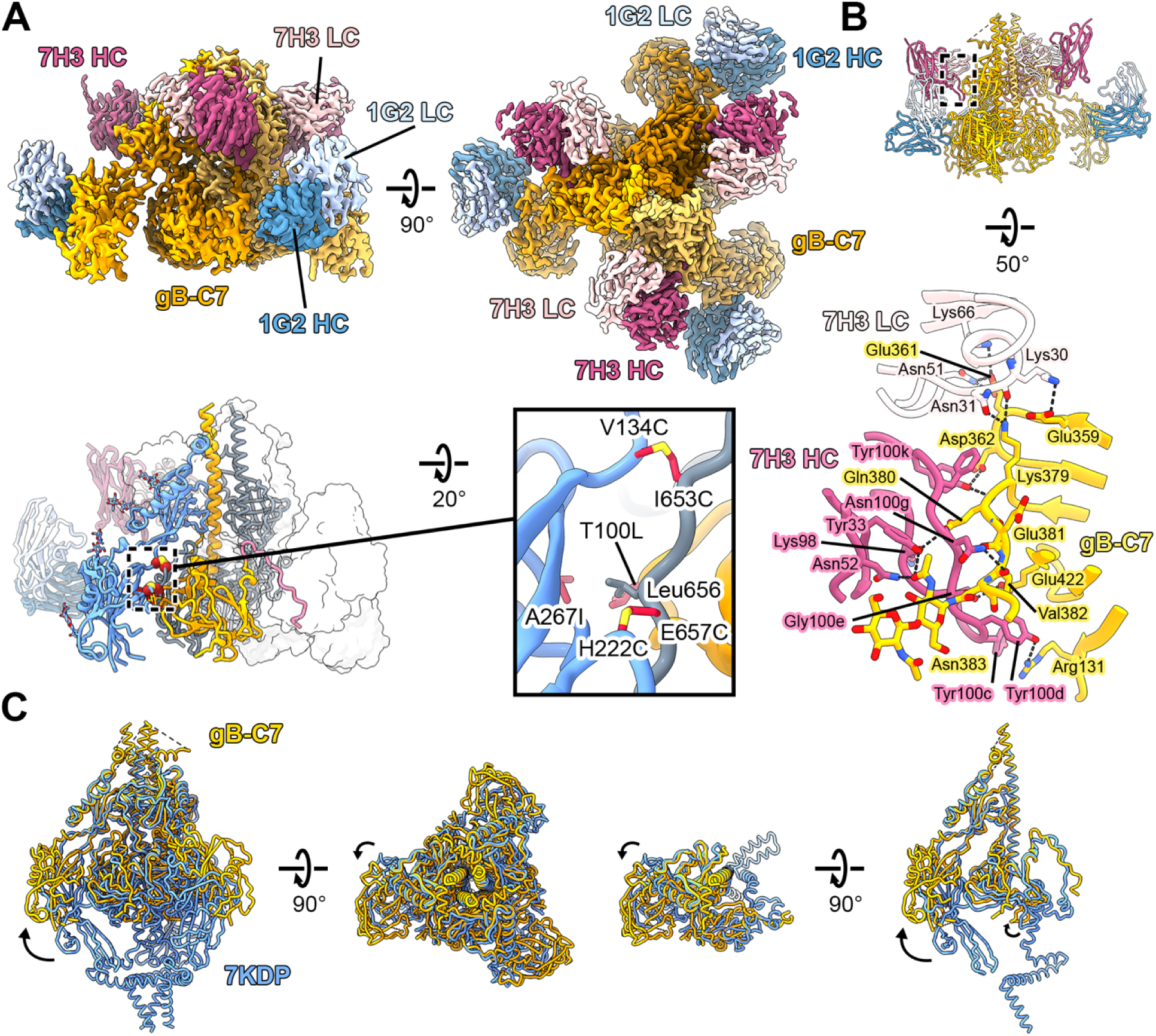
Cryo-EM structure of gB-C7 bound to 1G2 and 7H3 Fabs. (**A**) Side (*top, left*) and top (*top, right*) views of the composite EM map (global plus local refinement) of HCMV gB-C7 complexed with 1G2 and 7H3 Fabs shown above the gB-C7 complex model (*bottom, left*). One protomer of the model is colored as in Figure 1A and shown as a ribbon diagram, the second is colored gray and shown as a cartoon tube trace of the α-carbon backbone, and the third is shown as a transparent surface. The inset (*bottom, right*) shows a zoomed view of the substitutions that comprise design gB-C7 with side chains shown as sticks. (**B**) Side view of HCMV gB-C7 bound to 1G2 and 7H3 Fabs shown as a ribbon diagram (*top*) above a zoomed view of the binding interface between 7H3 and gB-C7 (*bottom*). 7H3 heavy chain (HC) is pink, 7H3 light chain (LC) is pale pink, and gB-C7 is yellow. In the zoomed interface, key residues are shown as sticks and hydrogen bonds are shown as black dashed lines. (**C**) The structure of gB-C7 (yellow) is superimposed with the previously determined structure of prefusion HCMV gB (blue, PDB ID: 7KDP) (*23*), both shown as cartoon tube traces of the α-carbon backbones. Side and top views are shown for the superimposition of HCMV gB both as a trimer (*left*) and as a single protomer (*right*). Shifts in domain arrangement are highlighted with arrows.

These residues adopt extended α-helices and are in agreement with an AlphaFold2 (AF2) predicted model (**Figure S9**) (*50*). Consistent with the previously reported prefusion HCMV gB structure (*23*), N-terminal residues 1–78 and a portion of the apex (residues 448–473) are unresolved in our map, likely due to the intrinsic flexibility of these regions. This is also consistent with the AF2 model, in which these regions are predicted to be unstructured. However, unlike the previously reported structure, we were unable to resolve the fusion loops of DI (residues 149–163, 192–197, and 233–246) or the majority of DV (residues 662–704). This is likely due to the omission of the MPR in our soluble ectodomain construct, which appears to be critical for interacting with the fusion loops and stabilizing DI (*23*). Further, our structure reveals that in the absence of the MPR, DI is shifted outward relative to its more compact positioning in the full-length prefusion HCMV gB structure reported previously (**Figure 4C**). Thus, the MPR or a soluble mimic likely needs to be present to maintain a compact prefusion conformation of gB.

All stabilizing substitutions in gB-C7 were designed to maintain DI in its prefusion position. The two disulfide bonds (V134C/I653C and H222C/E657C) were designed to bridge DI and DV (**Figures 4A**, **S10**), whereas the cavity-filling substitutions T100L/A267I were designed to stabilize the interface between DI and DIV via hydrophobic packing. We observed good map features for all side chains at these sites of substitution (**Figure S10**), although the map was not contiguous between the cysteines of the H222C/E657C disulfide. This may indicate that the H222C/E657C disulfides formed incompletely under these conditions, or they may have suffered damage from the electron beam (*51*).

The 1G2 and 7H3 interfaces were well resolved in the local and global maps, respectively, revealing extensive hydrogen bond networks that are maintained in both the prefusion and postfusion conformations (**Figures 4B**, **S11**). In line with the previously determined 1G2-bound postfusion HCMV gB structure (PDB ID: 5C6T), 1G2 binds a hydrophobic patch on DI, which corresponds to AD-5 of HCMV gB (*24*) (**Figure S11**). Consistent with a previously reported ns-EM reconstruction of postfusion HCMV gB in complex with 7H3 Fab (*52*), 7H3 binds the same DII epitope as the high-affinity, neutralizing antibody SM5-1 (*32*), which corresponds to AD-4 of HCMV gB (*23, 53*). The structures of gB Base and gB-C7 reported here are the first high-resolution structures of HCMV gB in complex with 7H3 Fab, providing a detailed characterization of the gB:7H3 interface, which includes 13 hydrogen bonds and 808 Å^2^ of buried surface area on gB-C7 (**Figure 4B**). Like SM5-1, binding appears to be largely mediated by an unusually long heavy chain CDR3 (*23, 53*).

Unlike gB-C7, which exhibits full 1G2 and 7H3 Fab occupancy in the 2.8 Å cryo-EM structure, 7H3 Fab bound to only one of the three protomers of gB Base (**Figure S3**). Comparison between occupied and unoccupied protomers reveals that 7H3 Fab displaces a flexible loop between domains II and III (residues 466–475) that localizes to the apex in prefusion gB and the trunk in postfusion gB (**Figure S3B**). When 7H3 is not bound, 10 additional residues of this DII/DIII loop are resolved, packing against the unoccupied 7H3 epitope. In a recently determined structure of postfusion HCMV gB in complex with SM5-1 Fab (PDB ID: 7KDD), SM5-1 binding also appears to displace the DII/DIII loop we resolve in our gB Base EM map (*23*). Our structural studies suggest that 7H3 and the DII/DIII loop compete to occupy a similar position in the postfusion conformation, explaining the partial occupancy of 7H3 Fab when complexed with gB Base observed by ns-EM and cryo-EM (**Figures 3D**, **S3**, **S5**). This partial occupancy, which has not been recognized in binding studies or reported previously, occurred reproducibly in our structural studies and suggests that 7H3 occupancy could be used to evaluate the prefusion character of gB proteins.

### Stabilized prefusion-like gB and postfusion gB are similarly immunogenic in mice

To assess the immunogenicity of the stabilized prefusion-like variants, three groups of eight BALB/c mice were immunized with gB antigens at weeks 0, 3, and 6. Each group was immunized with 2.5 µg of one of the following gB variants, which were selected to span a range of conformations: gB Base (postfusion), gB V134C/I653C (partially prefusion-stabilized), and gB-C7 (the most stabilized prefusion-like combination variant) (**Figures 5A**, **S12A**). To potentiate immune responses, the gB antigens were combined with CpG 1018 adjuvant plus alum (*54*). A fourth group of mice was immunized with gB-C7 without adjuvant to assess the impact of adjuvant on the quality of the immune response. Sera were collected 2 weeks after the third immunization (**Figure 5A**). To quantify antibody titers, we performed serum enzyme-linked immunosorbent assays (ELISAs) using antibody SM5-1 (*32*) expressed with murine IgG2a fragment crystallizable (mFc) domains as a standard. This neutralizing human antibody has a similar binding affinity for gB Base and gB-C7 (**Figure S12B**), consistent with previous findings that the SM5-1 Fab binds both postfusion and prefusion conformations (*23*). All gB variants were highly immunogenic, with serum antigen-binding IgG titers (1/ED_50_) ranging between 10^3^–10^6^ (**Figure S12C**, **D**). Antibodies elicited by immunization with adjuvanted gB Base, gB V134C/I653C, or gB-C7 exhibited similar binding to gB Base with SM5-1-binding-equivalent concentrations of 1.41, 1.34, and 1.14 mg/mL, respectively (**Figure 5B**, **Table S3**). Similar trends were observed when measuring antibody binding against gB-C7, where mice immunized with adjuvanted gB Base, gB V134C/I653C, or gB-C7 had similar responses, with SM5-1-binding-equivalent concentrations of 0.24, 0.22, and 0.21 mg/mL, respectively (**Figure 5C**). Predictably, mice immunized with gB-C7 without an adjuvant had significantly lower antibody titers against gB Base (0.07 mg/mL) and gB-C7 (0.01 mg/mL), indicating the immune response to gB-C7 can be enhanced by an adjuvant. Interestingly, immunization with postfusion or stabilized prefusion-like gB did not appear to bias antibody binding towards specific gB conformations (**Figure 5B, C**) and anti-gB Base titers correlated strongly with anti-gB-C7 titers (**Figure S13A**). These data are consistent with previous observations that many, if not all, known immunogenic gB epitopes are shared between conformations (*23*). Antibody binding to gB-C7 was lower for all treatment groups compared to gB Base (**Figure 5B, C**), suggesting there may be reduced availability of some prefusion epitopes. This is consistent with the differential exposure of certain regions of gB in the two conformations (*23*), including DIV and DV, which correspond to AD-1 and AD-6, respectively (*34, 55*) (*32, 33*).

**Figure 5.**
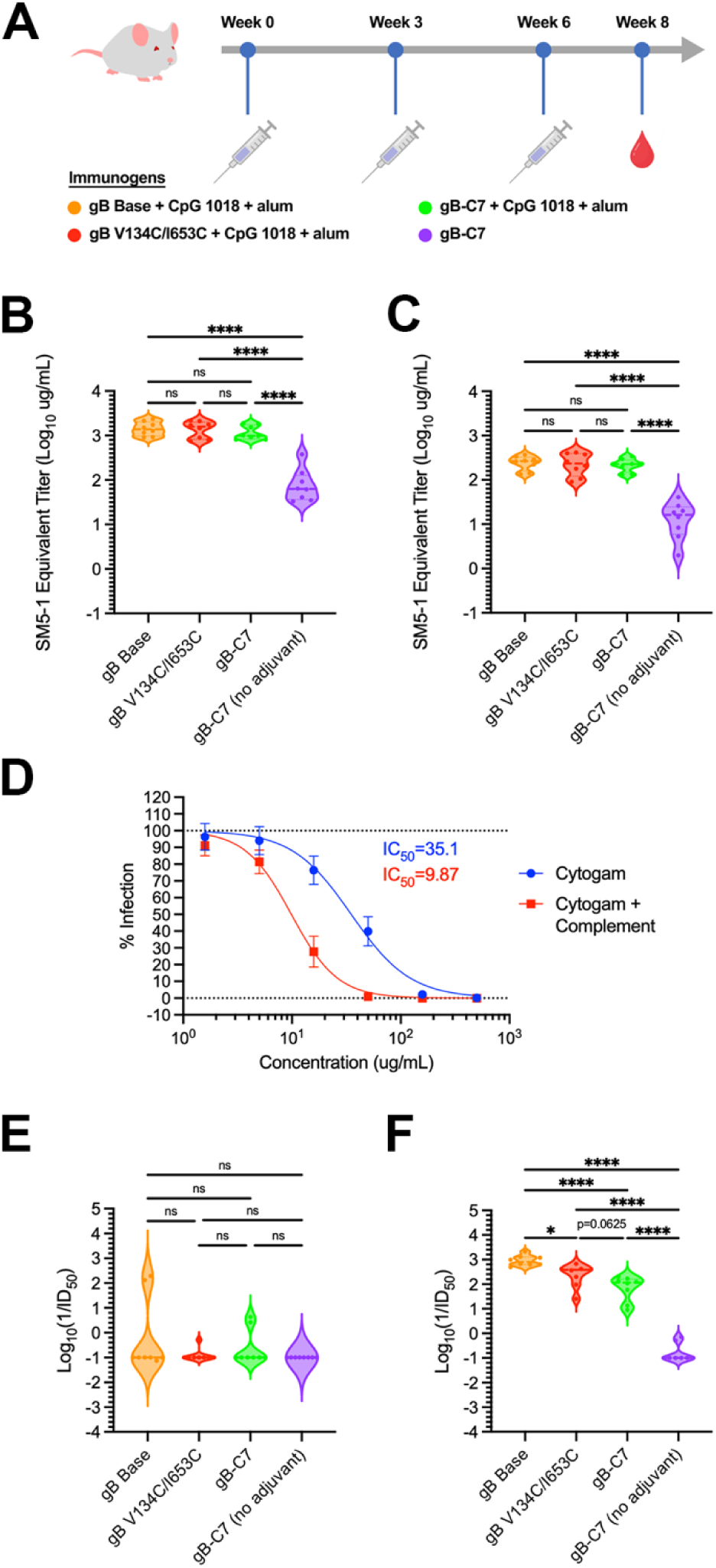
Engineered gB variants are immunogenic in mice. (**A**) Schematic of mouse immunization. 6– 8-week-old BALB/c mice (*n* = 8/group) were immunized at weeks 0, 3, and 6 with 2.5 μg of gB Base (orange), gB V134C/I653C (red), or gB-C7 (green), all adjuvanted with CpG 1018 plus alum or un-adjuvanted gB-C7 (purple). Blood samples were collected from mice 2 weeks after the week 6 injection. SM5-1-equivalent antibody concentrations of immunized mouse sera that bind to (**B**) gB Base or (**C**) gB-C7. Plots represent ELISA measurements relative to an SM5-1 mFc standard from two experimental replicates. AD169-GFP infection of human MRC-5 fibroblasts. (**D**) Neutralization by Cytogam is enhanced 3.6-fold in the presence of 12.5% guinea pig complement. Data is averaged across 16 plates with standard deviation shown. Neutralization by mouse sera in the (**E**) absence and (**F**) presence of 12.5% guinea pig complement. Plots represent averages from two independent experiments. In violin plots, horizontal lines represent the first quartile, median, and third quartile. Statistical significance was determined by one-way ANOVA in GraphPad Prism: * = p ≤ 0.05, ** = p ≤ 0.01, *** = p ≤ 0.001, **** = p ≤ 0.0001.

### Prefusion-like gB elicits weakly neutralizing, complement-enhanced antibodies

To test the hypothesis that immunization with prefusion-stabilized gB would produce superior neutralization titers, as is the case for multiple class I fusion proteins (*40, 56–58*), we measured the capacity of immunized mouse sera (with or without guinea pig complement) to neutralize HCMV laboratory strain AD169 infection of MRC-5 fibroblasts. We used Cytogam, a human hyper-immune globulin containing high titers of anti-HCMV polyclonal antibodies, as our positive control (*59*). In contrast to previous reports (*60*), but consistent with observations that complement can play a key role in neutralization by anti-gB monoclonal antibodies (*61*), we found that Cytogam-mediated neutralization of HCMV AD169 was enhanced 3.6-fold by the addition of 12.5% exogenous guinea pig complement (**Figure 5D**). The immunized mouse sera displayed weak neutralizing activity in the absence of complement (**Figure 5E**) and the addition of guinea pig complement also enhanced neutralization activity of sera from mice immunized with adjuvanted antigens (**Figure 5F**). No gB variants in this study consistently elicited complement-independent neutralizing antibodies in mice, although sera from several mice from different treatment groups exhibited measurable neutralization activity in the absence of complement (**Figures 5E**, **S12E**, **Table S3**). Notably, the two mice with the highest complement-independent responses were immunized with gB Base. Prior gB vaccination studies in humans (*62*) and rabbits (*63*) have also shown poor complement-independent neutralization of HCMV despite high anti-gB titers, consistent with the observation that gB vaccination often stimulates antibody responses directed toward non-neutralizing epitopes (*64*). Despite similar anti-gB binding titers across gB variants, we observed weaker complement-dependent neutralization responses against HCMV AD169 in fibroblasts from immunization with increasingly prefusion-like gB variants: gB Base appeared to be the most potent, followed by gB V134C/I653C and gB-C7 (**Figures 5F**, **S12**, **Table S3**). Anti-gB antibody titers correlated positively with complement-dependent neutralization but not complement-independent neutralization (**Figure S13**), suggesting gB binding and opsonization may be key factors in complement-dependent neutralization. Both gB Base and gB-C7 bound similarly to a panel of HCMV gB AD-2 antibodies, although the AD-2 site 1 (AD-2S1) TRL345 unmutated common ancestor (UCA) antibody (*65*) bound better to gB-C7 than gB Base (**Figure S14**). Collectively, the immunization data with the constructs described here suggest that prefusion-stabilized HCMV gB proteins are not superior immunogens to postfusion gB for the elicitation of potent neutralizing antibodies, at least against HCMV laboratory strain AD169 in fibroblasts.

## Discussion

The stabilization of fusion proteins in their prefusion conformations has been shown to be a beneficial approach to optimizing subunit vaccine antigens against a variety of viruses (*43, 66*). This strategy works well when the fusion protein contains neutralization-sensitive antigenic sites found exclusively on the prefusion conformation. This is true for the class I fusion protein respiratory syncytial virus (RSV) F, which displays two highly neutralization-sensitive sites exclusively at the apex of prefusion F (*67*). However, to the best of our knowledge, no class III fusion proteins have been sufficiently stabilized in their prefusion conformations to enable testing for improved immunogenicity. Aiming to stabilize a soluble HCMV gB construct in its prefusion conformation, we engineered and biochemically characterized a base construct, gB Base, along with 24 single substitution variants and 7 combination variants (**Figures 1**–**3**). With ns-EM, we found that certain constructs appeared to be partially stabilized in the prefusion conformation (**Figures 3D**, **S5**). We determined a 2.8 Å resolution cryo-EM structure of our most promising combination variant, gB-C7, confirming that it is stabilized in a prefusion-like conformation. We then immunized mice with gB Base, gB V134C/I653C (a single substitution variant), and gB-C7, and found that immunization with prefusion-like gB did not elicit a superior immune response as measured by the neutralizing antibody response against HCMV AD169 in fibroblasts (**Figure 5**).

Following the recently proposed model of HCMV gB rearrangement in which DI and DII are the first domains to undergo rearrangement from prefusion to postfusion (*23*), we hypothesized that substitutions aiming to tether DI in its prefusion position might globally maintain gB in the prefusion conformation (**Figure 1A**). In line with this hypothesis, single substitutions to DI displayed better expression and thermostability relative to gB Base and other domain substitutions (**Figures 1C, 2**). The membrane-distal apex of HCMV gB (DII and DIII) was stabilized in the prefusion conformation in the gB-C7 structure— in which all stabilizing substitutions were made to DI, DIV, and DV— demonstrating that stabilizing DI positioning can indeed stabilize gB in a prefusion-like conformation. However, compared to the recently published structure of wild-type HCMV gB in the prefusion conformation (PDB ID: 7KDP) (*23*), DI in the gB-C7 structure is shifted distally and only partially resolved by local refinement (**Figure 4C**), suggesting that DI is highly flexible and primed to rearrange in the absence of the MPR. Similar domain flexibility is displayed by the prefusion conformation of VSV G, a fellow class III fusion protein, that, in contrast to HCMV gB, does not possess an MPR and instead anchors its fusion loops directly into the viral membrane (*31, 68*). In comparison to the first crystal structure determined for VSV G (*31*), a second crystal structure published more recently displayed an 11° tilt of the fusion domain relative to the rest of the protein (*68*), underscoring the flexibility of this domain in ectodomain constructs of class III fusion proteins.

In designing single substitutions aimed at stabilizing HCMV in the prefusion conformation, we hypothesized that structure-based vaccine design strategies that have been successfully applied to class I fusion proteins would translate to class III fusion proteins. For severe acute respiratory syndrome coronavirus 2 (SARS-CoV-2) and Middle East respiratory syndrome coronavirus (MERS-CoV) spikes as well as RSV F and many other class I fusion proteins, proline substitutions—particularly helix capping proline substitutions—have been an effective method to stabilize the prefusion conformation (*56, 58, 69*). These substitutions generally target helix-turn-helix motifs in the prefusion conformation that rearrange to form extended helices in the postfusion conformation—by introducing prolines at these turns, the extended coiled-coil of the postfusion conformation is disfavored (*43*). Recently, proline substitutions aimed at capping a helix in DIII of HSV-1 gB and varicella-zoster virus (VZV) gB were shown to bias gB toward a prefusion-like conformation in the membrane (*30, 70*). However, the proline substitution was not sufficient to maintain the HSV-1 gB prefusion conformation in an ectodomain construct (*30*). Considering the recently published structure of prefusion HCMV gB (PDB ID: 7KDP) (*23*), as well as our structure of gB-C7, we hypothesize that a similarly located proline substitution in HCMV gB may distort the native architecture of the membrane-distal apex of the prefusion conformation. Of the five proline single substitution variants we tested, V645P, D646P, and L484P achieved around 50% increases in expression (**Figure 1C**). While L484P likely disrupts the central helix, V645P and D646P localize to a coil between α-helices in the prefusion conformation (**Figures 1B**, 4). This coil rearranges to an extended form in the postfusion conformation that packs between the central helices of DIII, a conserved motif seen in many postfusion herpesvirus fusion proteins (**Figures S1**, **S3**) (*24–27*). As such, proline substitutions targeted to this region may translate well to gB homologs. Overall, as most class III fusion proteins do not possess a prefusion helix-turn-helix motif that rearranges to an extended coiled-coil in the postfusion conformation, the strategy of proline capping to maintain the prefusion conformation may have limited potential in the context of herpesvirus fusion proteins. However, well-placed proline substitutions, such as V645P and D646P, may still have a positive impact on expression and stability.

We found that interprotomer disulfide substitutions, particularly those designed to bond DI to DV (H222C/E657C, N220C/E657C, and V134C/I653C), were the most effective method to stabilize HCMV gB in a prefusion-like conformation (**Figures 1C**, **3D**, **S5**, **4A**). As both DI and DV undergo conformational changes, these disulfide bonds stabilize the prefusion conformation by preventing rearrangement of the two domains. Interprotomer disulfide bonds have been used to stabilize several class I fusion proteins (*57, 71–74*), but often these disulfides reduce protein expression. However, here we found that interprotomer disulfide substitutions in HCMV gB generally had a neutral-to-positive impact on expression (**Figure 1C**) in addition to increasing thermostability and stabilizing gB in a prefusion-like conformation (**Figures 2C**, **3C**, **S5**, **4A**). As such, targeted interprotomer disulfide substitutions may be a general approach to stabilizing class III viral fusion proteins.

Since a vaccine candidate consisting of recombinant, structurally undefined HCMV gB paired with MF59 adjuvant achieved ∼50% efficacy in preventing HCMV infection in phase II clinical trials (NCT00125502, NCT00133497) (*16, 17*), development of a prefusion-stabilized HCMV gB antigen holds the potential to produce a more efficacious HCMV vaccine candidate. However, in our mouse immunization study, we found that gB-C7 was similarly immunogenic to gB Base (**Figure 5B, C**), and did not elicit higher titers of antibodies capable of neutralizing HCMV AD169 in fibroblasts. In fact, neutralization by sera from mice immunized with gB-C7 was dampened in comparison to gB Base (**Figure 5E, F**). There are several possibilities for the observed result: 1) insufficient stabilization of the prefusion-like gB construct, 2) the oligomerization of gB Base (**Figure S5**) may increase its immunogenicity, and 3) prefusion gB naturally lacks neutralization-sensitive epitopes found exclusively in this conformation, supported by the lack of prefusion-specific gB antibodies isolated to date. Indeed, as HCMV gB conformational rearrangement seems to be largely dominated by rigid body movements rather than reorganization of secondary structural elements, shared epitopes between prefusion and postfusion predominate (*23*). Nevertheless, the differing domain arrangement between prefusion and postfusion conformations suggests that it might be feasible to elicit antibodies that bind across domain interfaces to target potentially vulnerable, prefusion-specific epitopes. In future studies, our stabilized prefusion-like soluble HCMV gB-C7 immunogen should be assessed for its ability to elicit neutralizing antibodies against more clinically relevant HCMV isolates (*75*) and cell types (epithelial, endothelial cells) (*76, 77*). Further, non-neutralizing antibody functions that are associated with gB vaccine efficacy (*62, 78*) and decreased risk of congenital CMV transmission (*79, 80*) should be explored. HCMV gB-C7 should also facilitate isolation of prefusion-specific antibodies, particularly those that target regions at the well-stabilized membrane-distal apex, which may prove useful for therapeutic applications and as reagents for vaccine research and development.

## Methods

### Design scheme for prefusion-stabilized HCMV gB variants

The HCMV gB base construct (gB Base) comprises ectodomain residues 1–704 of HCMV gB Towne Strain (UniProtKB: P13201) with serine substituted at residues 246, 457, and 460 as previously described (*24*), followed by a C-terminal T4 fibritin (foldon) trimerization motif, an HRV3C protease recognition site, an octa-histidine tag, and a Twin-Strep affinity tag cloned into the mammalian expression vector pαH (*81*) and verified by DNA sequencing (**Figure 1A**). All HCMV gB variants were constructed into this plasmid by Gibson assembly and verified by DNA sequencing. Based on the HCMV gB prefusion structure (PDB ID: 7KDP) and postfusion structures (PDB IDs: 5CXF, 5C6T, 7KDD) (*23*) (*24, 25*), residues were considered for disulfide bond, polar, cavity-filling, and proline designs. Combinations were chosen to test whether pairing of designs could result in additive effects.

### Quantification of HCMV gB in HEK293F culture medium by bio-layer interferometry

Freestyle 293 cells were pipetted into vented 50 mL culture tubes at a density of 1 x 10^6^ cells/mL. Cultures were transiently transfected with 2 μg of plasmids encoding HCMV gB variants using polyethyleneimine (PEI) as a transfection reagent (25,000 MW at a ratio of 9:1 PEI to DNA). Transfected cultures were maintained in a shaking incubator at 175 RPM (37 °C, 8% CO_2_) for 6 days. Culture medium was harvested by centrifugation at 1,000 x *g* to pellet cells. The supernatant was then transferred to microcentrifuge tubes and centrifuged at high speed (20,000 x *g*) to pellet insoluble material. The clarified medium was diluted 5-fold with 1X HBS EP+ (10 mM HEPES pH 8.0, 150 mM NaCl, 3 mM EDTA, 0.005% (w/v) Tween 20, 0.01% (w/v) sodium azide), then pipetted into black-walled 96-well plates (Greiner Bio-One) and loaded into a bio-layer interferometer (Octet RED96, ForteBio). Anti-human IgG Fc capture (AHC) biosensor tips (Sartorius) were equilibrated for ten minutes in a 1:4 mixture of conditioned Freestyle 293 medium and 1X HBS EP+ to match the diluted media, then dipped into anti-foldon IgG MF5, which was a gift from Dr. Vicente Mas (Instituto de Salud Carlos III), followed by HCMV gB medium. The amount of HCMV gB in each sample of clarified medium was quantified by fitting the response as a function of time to a straight line over the linear portion of the binding curve. We used a dilution series of purified HCMV gB in conditioned HEK293F medium (10 mg/mL to 0.156 mg/mL) to generate a standard curve, which allowed us to report the absolute concentration of HCMV gB in the medium. Data were plotted as an average of three independent biological replicates using GraphPad Prism 9.5.1 and subsequent versions (GraphPad Software).

### Purification of HCMV gB by affinity and size-exclusion chromatography

HEK293F cell cultures (40 mL) were grown to a density of 1 x 10^6^ cells/mL and then transiently transfected with 25 μg (9:1 PEI to DNA ratio). After 5–6 days, medium was harvested by centrifugation, 0.22 µm filtered, then passed over Strep-Tactin Sepharose resin (IBA Lifesciences) by gravity, washed with 3 column volumes of 1X PBS, and eluted with Strep-Tactin elution buffer (100 mM Tris-Cl pH 8.0, 150 mM NaCl, 1 mM EDTA, and 2.5 mM desthiobiotin) (IBA Lifesciences). Elution fractions were analyzed by SDS-PAGE. Fractions containing HCMV gB were pooled, concentrated with Amicon Ultra centrifugal filters (MilliporeSigma), and flash-frozen in liquid nitrogen. Samples were thawed in a room temperature water bath just before injection onto a Superose 6 Increase 10/300 GL column (Cytiva). The running buffer was composed of 2 mM Tris pH 8.0, 200 mM NaCl, and 0.02% (w/v) sodium azide. Desired fractions were pooled, concentrated, aliquoted, and flash-frozen in liquid nitrogen for further analysis.

### Differential scanning fluorimetry

Concentrated, purified HCMV gB samples were diluted to 0.6 mg/mL and mixed with dilute SYPRO Orange. We used a concentrated SYPRO Orange stock (ThermoFisher Scientific, 5,000X SYPRO Orange Protein Gel Stain in DMSO), which was diluted ∼83 fold in 2 mM Tris pH 8.0, 200 mM NaCl, and 0.02% (w/v) sodium azide to obtain a working solution. The absorbance of the working SYPRO Orange solution was 0.295 at 468 nm (1 mm path length in a NanoDrop One, ThermoFisher Scientific). For each sample, 15 μL of 0.6 mg/mL HCMV gB was mixed with 5 μL of dilute SYPRO Orange working solution in white-walled 96-well PCR plates (VWR), which were then sealed and loaded into a LightCycler 480 differential scanning fluorimeter (Roche) equipped with a 100W xenon excitation lamp. We used a 465 nm excitation filter (25 nm half band width) and a 580 nm emission filter (20 nm half band width). The fluorescence in each well was monitored as the temperature of the plate was increased from 25 °C to 90 °C. Data were plotted as the change in fluorescence with respect to temperature (dF/dT) as a function of temperature using GraphPad Prism 9.5.1 and subsequent versions (GraphPad Software).

### Expression and purification of Fabs for structural and biophysical studies

For 1G2 and 7H3, a stop codon was introduced before the hinge region of the heavy chains to generate fragments of antigen binding (Fabs). HEK293F cell cultures were grown to a density of 1 x 10^6^ cells/mL and then transiently transfected with plasmids encoding the heavy and light chains at a 1:1 ratio (9:1 PEI to DNA ratio). Cultures were harvested 6 days after transfection and the medium was separated from the cells by centrifugation. Supernatant was passed through a 0.22 μm filter and then concentrated using a tangential flow filtration cassette (PALL) before Fab was purified with CaptureSelect™ IgG-CH1 Affinity Matrix (ThermoFisher). Bound Fab was eluted in a buffer containing 100 mM glycine pH 3.0. The protein elution was immediately neutralized with 1 M Tris pH 8.0, and then further purified by size-exclusion chromatography using a HiLoad 16/600 Superdex 200 pg column (GE Healthcare) into 2 mM Tris pH 8.0, 200 mM NaCl, and 0.02% (w/v) sodium azide. All protein samples were concentrated using Amicon Ultra centrifugal filter units (MilliporeSigma) to between 1 and 10 mg/ml, then flash-frozen in liquid nitrogen and stored at −80 °C.

### Negative-stain electron microscopy

Concentrated, purified HCMV gB constructs were mixed with Fabs (1G2, 7H3, or both 1G2 and 7H3) at a ratio of 1.2:1 (Fab:gB), incubated for 15–30 minutes at room temperature, then diluted to a working concentration of 0.02–0.1 mg/mL in a buffer composed of 2 mM Tris pH 8.0, 200 mM NaCl, and 0.02% (w/v) sodium azide. Samples diluted at the working concentration were immediately applied to glow-discharged copper-supported carbon grids (Formvar, 400 mesh), incubated for 30 seconds, washed three times with buffer composed of 2 mM Tris pH 8.0, 200 mM NaCl, and 0.02% (w/v) sodium azide, then stained with 2% methylamine tungstate (Nano-W, Nanoprobes) and air dried. Grids were loaded onto one of two transmission electron microscopes (TEMs): (i) a Japan Electron Optics Laboratory (JEOL) 2010F TEM or (ii) a JEOL NEOARM. The nominal magnifications for the 2010F and NEOARM were 60,000X (pixel size = 3.6 Å) and 50,000X (pixel size = 2.16 Å), respectively. Both microscopes operated at 200 kV and were equipped with OneView cameras (Gatan). Micrographs were acquired in 2k x 2k mode for the 2010F and 4k x 4k mode for the NEOARM using Digital Micrograph (Gatan), then exported to cryoSPARC v4 (Structura Biotechnology) for contrast transfer function (CTF) correction, particle picking, and 2D classification (*82*). 3D volumes were generated using ab initio reconstruction and data were further processed through heterogeneous and homogenous refinements. Structural figures were produced using ChimeraX (*83*).

### Production of HCMV gB protein for cryo-EM

FreeStyle 293 F cells were grown to a density of 1 x 10^6^ cells/mL and transiently transfected with plasmids encoding either HCMV gB Base or gB-C7 with PEI (9:1 PEI to DNA ratio). Approximately 4 hours after transfection, 5 mM kifunensine was added to the cell media. After 5–6 days, medium was harvested by centrifugation and 0.22 µm filtered. Secreted proteins were concentrated and exchanged into PBS using a tangential flow filtration cassette (PALL), then purified by Strep-Tactin Sepharose resin (IBA Lifesciences), washed with 5 column volumes of 1X PBS, and eluted with Strep-Tactin elution buffer (100 mM Tris-Cl pH 8.0, 150 mM NaCl, 1 mM EDTA, and 2.5 mM desthiobiotin) (IBA Lifesciences). Eluate was concentrated with Amicon Ultra centrifugal filter units (MilliporeSigma). For gB-C7, eluate was then treated with 5% (w/w) HRV3C protease at 4 °C overnight to remove affinity tags. For gB Base, eluate was flash-frozen in liquid nitrogen. Samples of gB Base were thawed in a room temperature water bath just before injection onto a HiLoad Superose 6 16/70 pg column (Cytiva). For gB-C7, solution was injected onto a HiLoad Superose 6 16/70 pg column (Cytivia) following HRV3C digestion. The running buffer was composed of 2 mM Tris pH 8.0, 200 mM NaCl, and 0.02% (w/v) sodium azide. Desired fractions were pooled, concentrated, aliquoted and flash-frozen in liquid nitrogen for further analysis.

### Cryo-EM sample preparation and data collection

All cryo-EM data sets were collected on an FEI Titan Krios operating at 300 kV and equipped with a K3 direct electron detector (Gatan) (Table S1). CF-400 1.2/1.3 grids (Electron Microscopy Sciences) were glow discharged for 60 seconds at 15 mAmps (PELCO easiGlow™ Glow Discharge Cleaning System).

A 3.0 mg/mL solution of gB Base complex was prepared by incubating roughly equimolar concentrations of gB Base and 1G2 Fab and a 2-fold molar excess of 7H3 Fab in 2 mM Tris pH 8.0, 200 mM NaCl, 0.02% (w/v) sodium azide, 3% (v/v) glycerol, and 0.01% (w/v) amphipol A8-35. The complex was incubated for 30 minutes at 4 ℃ before adding 10X CMC CHAPS (VitroEase™ Buffer Screening Kit, ThermoFisher Scientific) to a final concentration of 0.5X CMC. Immediately after, 3 µL of the gB Base complex solution was applied onto grids. Grids were double blotted and subsequently plunge-frozen in liquid ethane using a Vitrobot Mark IV (ThermoFisher Scientific) set to 100% humidity and 4 ℃ with a blot time of 5 seconds, a blot force of -1, and a wait time of 5 seconds. Data were collected at 105,000x magnification corresponding to a calibrated pixel size of 0.83 Å. A total of 8,679 exposures were collected for the gB Base complex data set, 1,308 exposures of which were collected without tilt and 7,371 exposures of which were collected with a 30° tilt. Data were collected using SerialEM 3.9.0 beta (*84*).

A 4.0 mg/mL solution of gB-C7 complex was prepared by incubating roughly equimolar concentrations of gB-C7, 1G2 Fab, and 7H3 Fab in 2 mM Tris pH 8.0, 200 mM NaCl, 0.02% (w/v) sodium azide, 3% (v/v) glycerol, and 0.01% (w/v) amphipol A8-35. The complex was incubated for 30 minutes at 4 ℃ before adding 10X CMC CHAPS (VitroEase™ Buffer Screening Kit, ThermoFisher Scientific) to a final concentration of 0.25X CMC. Immediately after, 3 µL of the gB-C7 complex solution was applied onto grids. Grids were blotted and subsequently plunge-frozen in liquid ethane using a Vitrobot Mark IV (ThermoFisher Scientific) set to 100% humidity and 4 ℃ with a blot time of 5 seconds, a blot force of -1, and a wait time of 5 seconds. Data were collected at 105,000x magnification corresponding to a calibrated pixel size of 0.83 Å. A total of 12,524 exposures were collected without tilt using SerialEM 3.9.0 beta.

### Cryo-EM data processing, model building, and refinement

The data processing began with gain reference correction before the micrographs were imported into cryoSPARC Live. All data sets were initially processed in cryoSPARC Live and subsequently processed in cryoSPARC v4.2.0 and subsequent versions. Motion correction, patch CTF estimation, defocus estimation, micrograph curation, blob particle picking, particle extraction, and 2D classification were initially performed in cryoSPARC Live for all data sets. Micrographs and picked particles were then exported into CryoSPARC, and particle curation through iterative rounds of 2D classification was performed using cryoSPARC v4.2.0 and subsequent versions. 3D volumes were generated using ab initio reconstruction, and data were further processed through heterogeneous refinement, homogenous refinement, and subsequent nonuniform homogeneous refinement of final classes. For the gB-C7 complex, a mask was created around domain I and 1G2 using ChimeraX and imported to cryoSPARC for local refinement. The resulting local map was combined with the global map using PHENIX combine_focused_maps (*49*). The EM processing pipelines for the gB Base and gB-C7 data sets are summarized in Figures S1 and S3, respectively. Initial models of gB Base, 1G2 Fab, and 7H3 Fab were predicted using AlphaFold2 (AF2) (*50*) while an initial model of gB-C7 was built with ModelAngelo (*85*). Models were then fit into the experimental cryo-EM maps in ChimeraX (*83*). Iterative model building and refinement were performed using PHENIX (*49*), COOT (*86*), and ISOLDE (*87*). The protein interfaces, surfaces, and assemblies (PISA) service at the European Bioinformatics Institute was used to determine buried surface area and interacting residues (*88*), and structural figures were produced using ChimeraX (*83*).

### Data processing and statistical analysis

Statistical analyses were performed using GraphPad Prism 9.5.1 and subsequent versions (GraphPad Software). We also used GraphPad Prism 9.5.1 and subsequent versions to plot the data. Information about the statistical tests performed can be found in the figure legends.

### Animal experiment

In vivo immunogenicity of gB protein in combination with adjuvants was evaluated in 6–8 week old BALB/c mice (Charles River) at Aragen Bioscience (Morgan Hill, CA). All procedures were carried out under institutional IACUC-approved protocols. Groups of 8 mice were immunized by the intramuscular route at Days 0, 21, and 42 with 2.5 mg/dose of gB protein (gB Base, gB V134C/1653C, or gB-C7) in the presence or absence (gB-C7 only) of CpG 1018 adjuvant (10 mg; Dynavax) and aluminum hydroxide (alum; Alhydrogel®; 50 mg; InvivoGen). Immunogens were prepared by first mixing gB protein with alum for 30 minutes, followed by addition of CpG 1018 for an additional 5 minutes of mixing. Mice were injected within 1 hour of immunogen preparations. Blood samples were collected by the submandibular route at Day 0 and Day 42 (post second immunization) and by cardiac puncture at Day 56 (day of sacrifice) for serum harvests.

### HCMV preparation

Human MRC-5 fibroblasts (ATCC, CCL-171) were cultured in Dulbecco’s modified Eagle medium supplemented with 10% FBS, 100 U/mL penicillin and 100 μg/mL streptomycin at 37 °C, 5% CO_2_. Virus stocks were produced as described from AD169-GFP BAC (*89*), which was a gift from Dr. Thomas Shenk (Princeton). In brief, MRC-5 cells were transfected with the BAC to generate a P0 stock. MRC-5 cells grown to 80% confluent monolayers were then infected using the P0 stock at an MOI of 0.01 and cultured for ∼2 weeks. Virus was concentrated from the supernatant using centrifugation with a 17% sorbitol cushion to generate the P1 virus used in all experiments. The titer of viral stocks was measured by infecting MRC-5 cells grown to 80% confluence on 24-well culture plates with serially diluted virus in a 500 μL volume. After 1 hour, cells were washed once with cell culture medium and incubated overnight. About 17 hours post-infection, cells were detached from the plate and %GFP positive cells were measured on an Attune flow cytometer (ThermoFisher Scientific). Titer was quantified by the following calculation and reported as infectious units (IU) per mL:

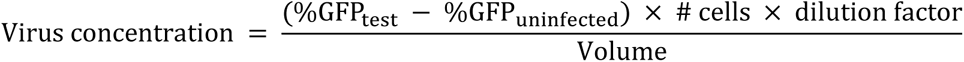

### Immunized serum ELISAs

SM5-1 antibody with murine IgG2a Fc (SM5-1 mFc) was produced in ExpiCHO cells via transient transfection and purified from cell culture supernatant by Protein A affinity chromatography for use as an ELISA standard. ELISAs were performed at room temperature using PBS + 0.1% (w/v) Tween 20 (PBST) as the wash buffer and 5% milk in PBST as the blocking buffer. All wells were washed three times with 300 μL PBST in between coating, blocking, primary, secondary, and developing steps. High-binding 96-well plates were coated with gB Base or gB-C7 proteins in Dulbecco’s PBS at a concentration of 1 μg/mL for 1 hour. Plates were then blocked with blocking buffer for 30 minutes. The standard (5 μg/mL SM5-1 mFc mixed with 1:100 unimmunized mouse serum) and 1:100 diluted immunized mouse sera were serial diluted by a factor of √10 in blocking buffer and applied to the plate for 1 hour. The secondary antibody used was goat anti-mouse Ig HRP (Southern Biotech, Cat. #1010-05) diluted 1:2000 in blocking buffer, which was applied to the plate for 1 hour. Plates were then developed using 3,3′,5,5′-tetramethylbenzidine (TMB) substrate solution and quenched with 1 N HCl. Absorbance at 450 nm was recorded using a SpectraMax M5 plate reader (Molecular Devices). ELISA curves were fit to a 4-parameter logistic (4PL) curve using Microsoft Excel and normalized to the standard of their respective plates. The concentrations of antibodies in immunized mouse sera relative to the standard were estimated by fitting 4PL curves of serum samples to the standard curve. Antibody titers are shown as averages of two technical replicates from two separate experimental replicates and were log-transformed, plotted, and evaluated for statistical differences between means in GraphPad Prism 9.5.1 and subsequent versions (GraphPad Software).

### In-vitro neutralization assay

One day prior to the neutralization assay, MRC-5 cells were seeded at 5–10 x 10^3^ cells/well in a cell-culture treated black, clear-bottom 96-well plate. Mouse sera were heat-inactivated (56 °C for 30 minutes) and √10 serially diluted (4 x 10^-2^–1.3 x 10^-4^). For assays including complement, guinea pig complement (Cedarlane, CL4051) was mixed with cell culture media prior to addition to virus and sera at a final concentration of 12.5%. Cytogam hyper-immune globulin was used as a positive control and was √10-fold serially diluted (100 to 0.32 μg/mL). Sera or antibody were mixed with 2500 IU AD169-GFP in culture medium and co-incubated with sera in a 50 µL volume for 2 hours at 37 °C, 5% CO_2_. Cells were co-incubated with virus/antibody mixture for 1 hour at 37 °C, 5% CO_2_. Cells were then washed, overlayed with 100 µL culture medium, and incubated for 17–20 hours at 37 °C, 5% CO_2_. Wells were then stained with 2 μM Hoechst 33342 nuclear stain (Invitrogen H1399) and imaged on a Cytation C10 confocal plate reader (BioTek). All experimental conditions were performed in duplicate wells. Uninfected and infected, no-antibody controls were performed in quadruplicate on every plate. Nuclei and GFP^+^ cells were then counted using Gen5 imaging software (v3.13, BioTek) and used to calculate the ratio of infected cells to number of nuclei (R). %Infection was calculated using the following equation:

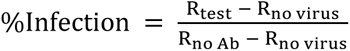

Averages of two experimental replicates were analyzed as ID_50_ and IC_50_ curves calculated by fitting to a 4PL curve (inhibitor, normalized response) in GraphPad Prism 9.5.1 and subsequent versions (GraphPad Software).

## Acknowledgments

We thank members of the Texas Materials Institute EM facility (Karalee Jarvis, Raluca Gearba, and Xun Zhan) and the UT Austin Center for Biomedical Research Support (Michelle Mikesh) for technical assistance with negative-stain EM. We thank members of the Sauer Structural Biology Lab (Evan Schwartz, Axel Brilot) for technical assistance with cryo-EM data collection. We thank Kaci Erwin and James Guerra for technical assistance with mammalian cell culture, and we thank Justine Meccio for administrative support. We thank Emily J. Rundlet for providing helpful comments on the manuscript.

## Funding

Funding for these studies was provided by Dynavax Technologies Corporation. The Sauer Structural Biology Laboratory is supported by the University of Texas College of Natural Sciences and by award RR160023 from CPRIT.

## Author contributions

Conceptualization, M.R.S., M.J.B, and J.S.M.; investigation, M.R.S., P.O.B., A.G.L., A.R.R., J.A.G., R.S.M., L.Z., N.V.J., C.-L.H., M.C., K.K., J.D.C., D.Y., and M.J.B.; visualization, M.R.S., P.O.B., A.G.L., K.K.; writing–original draft, M.R.S., P.O.B., A.G.L., and J.S.M.; writing–reviewing & editing, M.R.S., P.O.B., A.G.L., A.R.R., R.S.M., L.Z., N.V.J., C.-L.H., K.K., J.D.C., S.R.P., J.A.M., D.Y., M.J.B., and J.S.M.; supervision, J.D.C., S.A.P., J.A.M., D.Y., M.J.B., and J.S.M.

## Competing Interests

J.S.M., M.R.S., P.O.B., C.-L.H., L.Z., R.S.M., N.V.J., J.A.G., and A.R.R. are inventors on a patent application entitled *“Prefusion-Stabilized CMV gB Proteins”* (PCT/US2023/073369). J.D.C., D.Y., and M.J.B. are employees of Dynavax and hold Dynavax stock. The other authors declare that they have no competing interests.

## Data and materials availability

All data needed to evaluate the conclusions in the paper are present in the paper and/or the Supplementary Materials. EM density maps and structure models for prefusion and postfusion gB complexes are accessible through EMD-43667 and EMD-43670, 43671, and 43672 and PDB ID: 8VYM and 8VYN, respectively.

