## Supplemental Figures and Tables for "Structure-based design of a soluble human cytomegalovirus glycoprotein B antigen stabilized in a prefusion-like conformation"

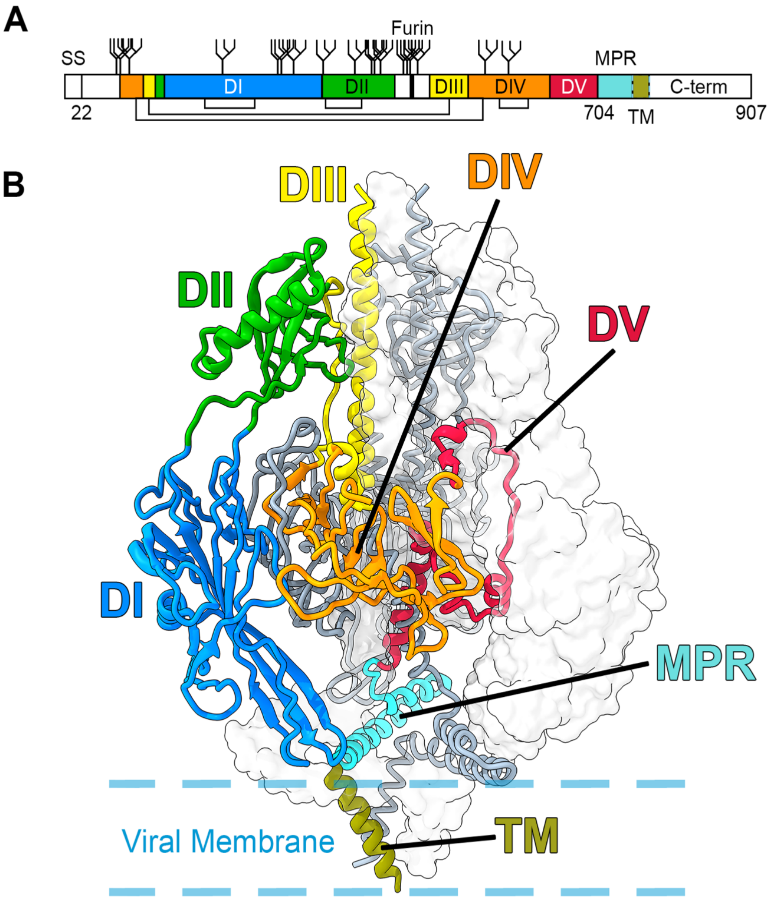


**Figure S1. Domain architecture of HCMV gB**. (**A**) Schematic of wild-type HCMV gB. Native disulfide bonds are shown as connected lines. N-linked glycosylation sites are shown as branched lines. The native furin cleavage site is shown as a thick black line. The N-terminal signal sequence (SS) is shown as a white box. Structural domain I (DI) is colored blue, DII is colored green, DIII is colored yellow, DIV is colored orange, DV is colored red, the membrane-proximal region (MPR) is colored cyan, the transmembrane domain (TM) is colored olive and flanked by dashed blue lines, and the C-terminal domain (C-term) is white. (**B**) Side view of trimeric prefusion HCMV gB (PDB ID: 7KDP) (*23*). One protomer is colored as in A and shown as a ribbon diagram, one protomer is colored gray and shown as a cartoon trace of the α-carbon backbone, and one protomer is shown as a transparent surface. The approximate location of the viral membrane is shown as dashed blue lines for orientation.

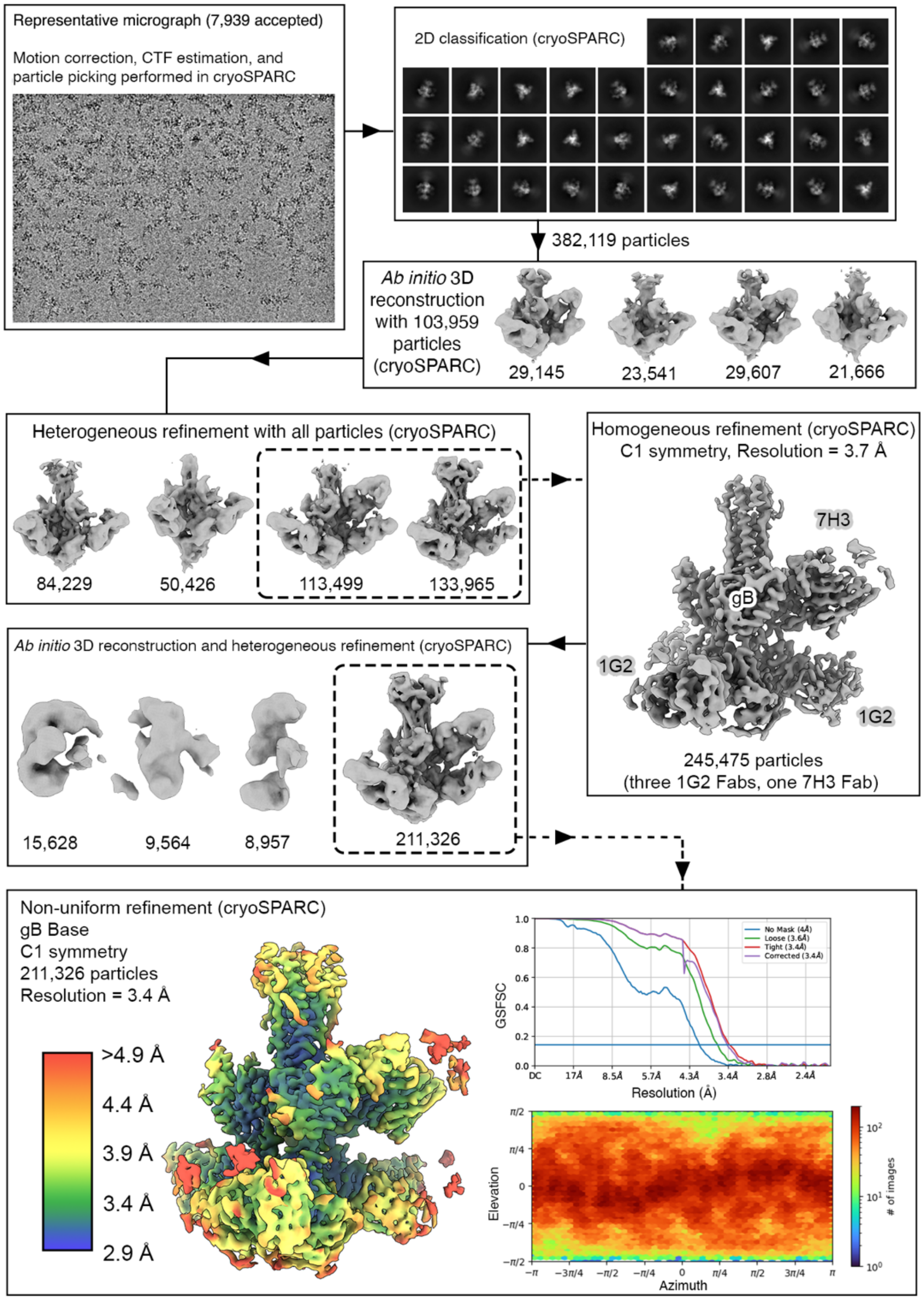


**Figure S2. Cryo-EM data collection and processing summary for the structure of gB Base in complex with 1G2 and 7H3 Fabs**. Cryo-EM processing workflow consisted of the following steps performed in cryoSPARC v4.2.0: motion correction, CTF estimation, particle picking, 2D classification, *ab initio* 3D reconstruction, iterative heterogenous refinement, homogenous refinement, and non-uniform refinement with C1 symmetry. The number of particles used in each step is listed. The final, highest resolution reconstruction is shown at the bottom of the figure, colored as a rainbow according to the estimated local resolution from 2.9 Å (blue) to 4.9 Å (red) resolution. Gold-standard Fourier shell correlation and directional distribution plots are shown to the right of the high-resolution reconstruction.

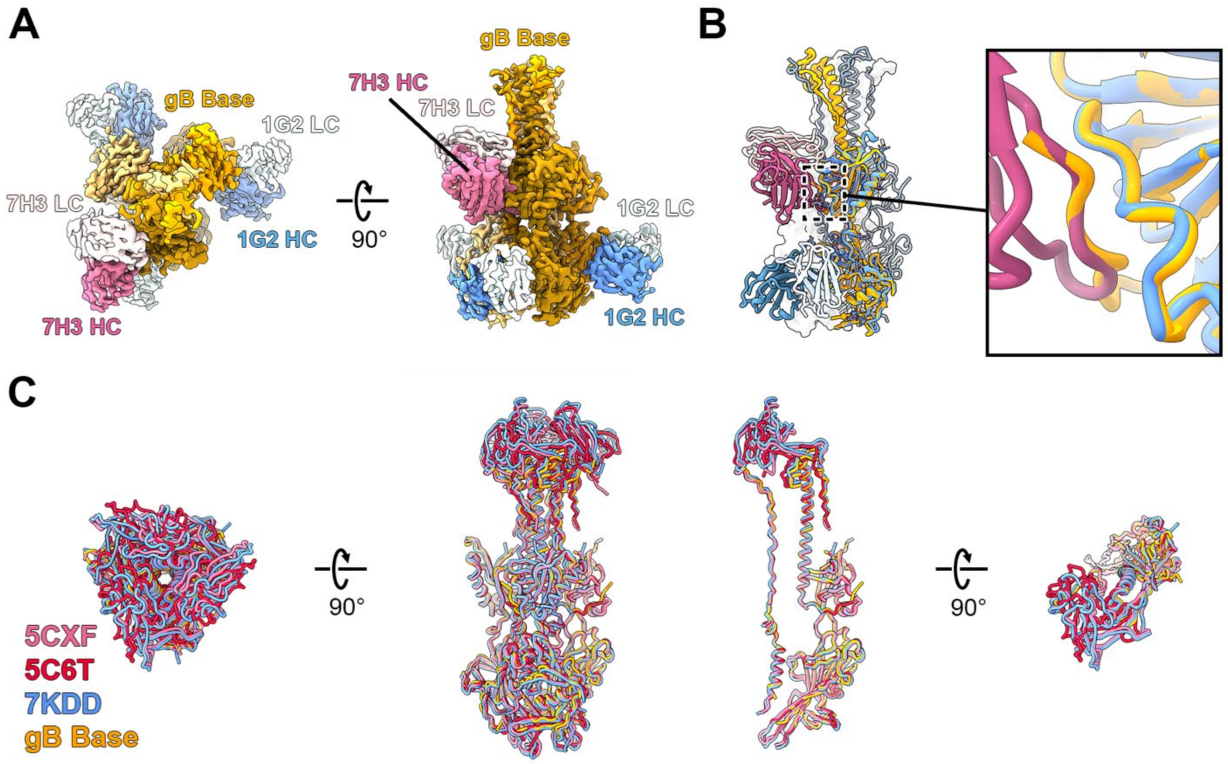


**Figure S3. Cryo-EM structure of gB Base bound to 1G2 and 7H3 Fabs**. (**A**) Top (*left*) and side (*right*) views of the EM map of HCMV gB ectodomain gB Base complexed with 1G2 and 7H3 Fabs. (**B**) The gB Base complex model. One protomer of the model is colored as in Figure 1A and shown as a ribbon diagram, the second is colored gray and shown as a cartoon tube trace of the α-carbon backbone, and the third is shown as a transparent surface. The structure of an unbound protomer (yellow) is shown as a ribbon diagram and superimposed with the protomer bound by 7H3 Fab. The inset shows a zoomed view of a flexible loop displaced by 7H3 Fab. (**C**) The structure of gB Base (yellow) is superimposed with the previously determined structures of postfusion HCMV gB (PDB IDs: 5CXF, 5C6T, and 7KDD in red, pink, and blue, respectively) (*23-25*), all shown as cartoon tube traces of the α-carbon backbones. Top and side views are shown for the superimposition of postfusion HCMV gB both as a trimer on the left and as a single protomer on the right.

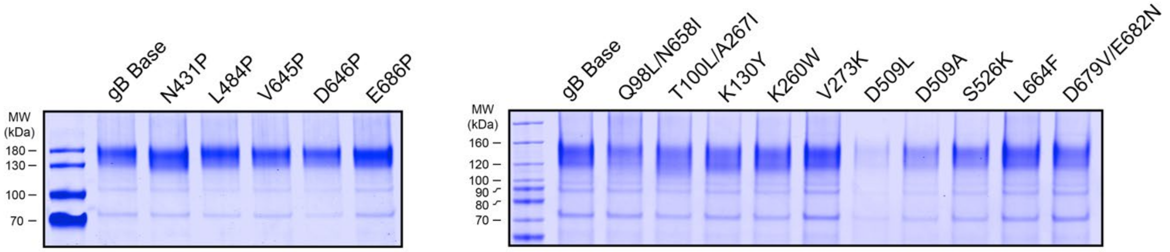


**Figure S4. Reducing SDS-PAGE for single and paired substitution variants**. The molecular weight ladder is in the left-most lane of each gel with standards indicated at the left in kDa.

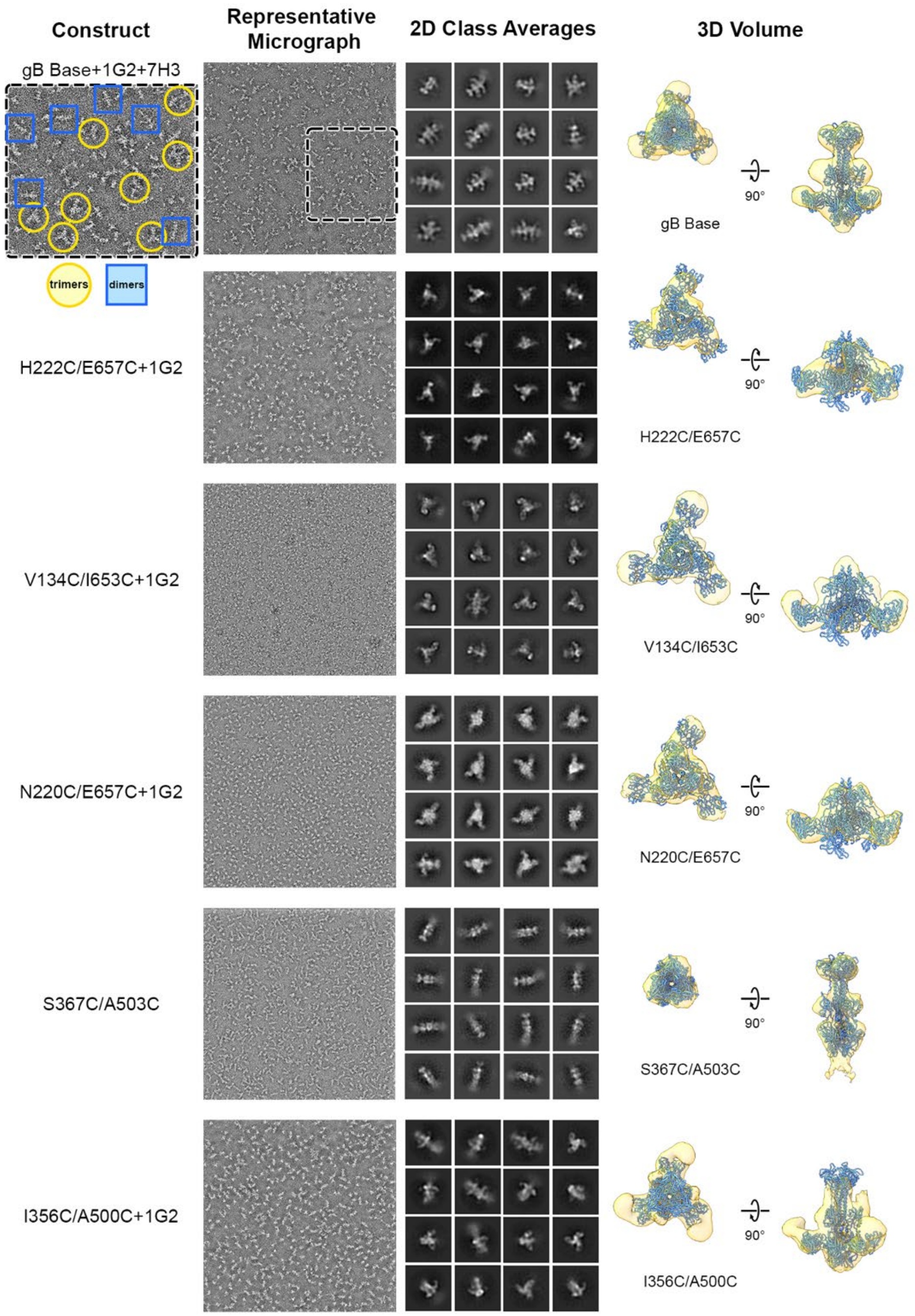


**Figure S5 (continues on next page). Negative-stain EM for gB Base and select gB variants**. Representative negative-stain EM (ns-EM) micrographs, 2D class averages, and low-resolution 3D reconstructions with imposed C3 symmetry for gB Base and select gB variants. The gB Base inset (*top, left*) shows a zoomed view of a representative micrograph with higher-order oligomers (rosettes) highlighted with blue squares and yellow circles, indicating dimers of gB trimer and trimers of gB trimer, respectively.

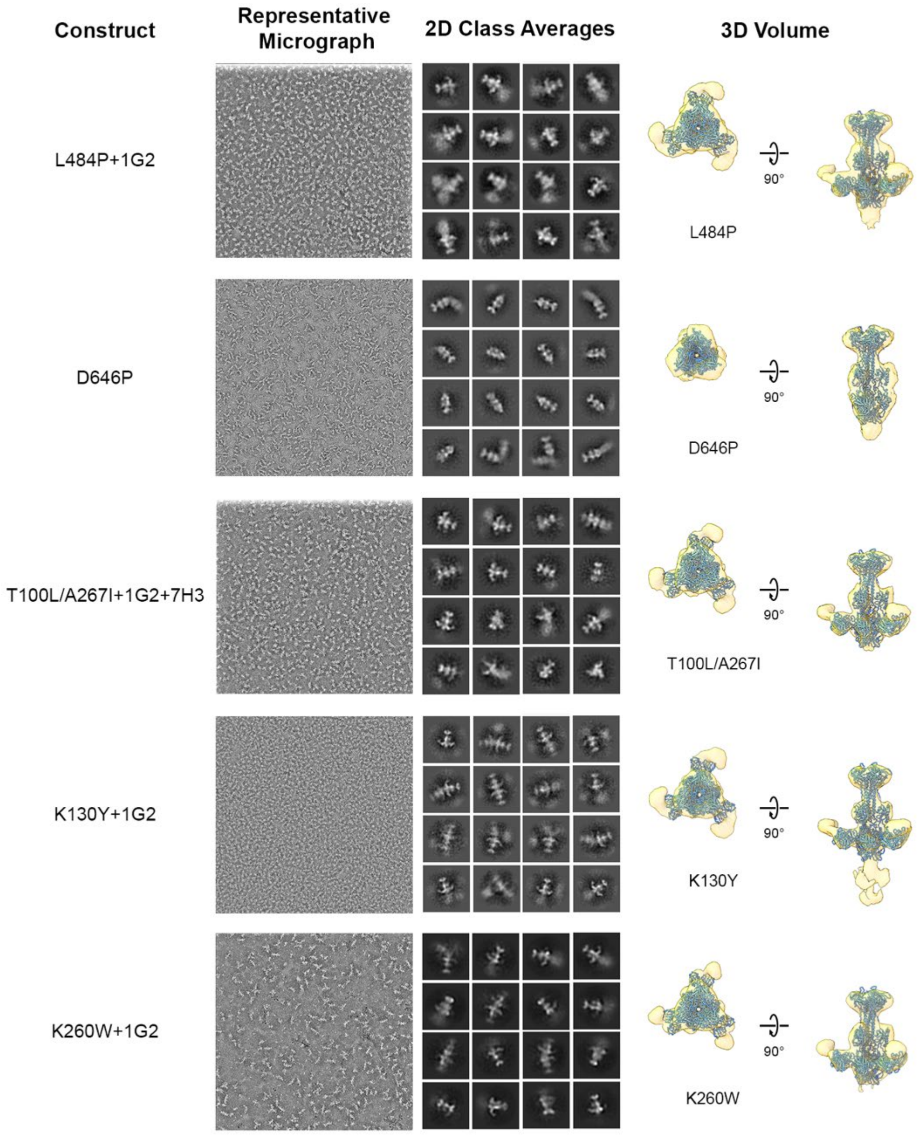


**Figure S5 (continued from previous page). Negative-stain EM for gB Base and select gB variants**. Representative negative-stain EM (ns-EM) micrographs, 2D class averages, and low-resolution 3D reconstructions with imposed C3 symmetry for gB Base and select gB variants.

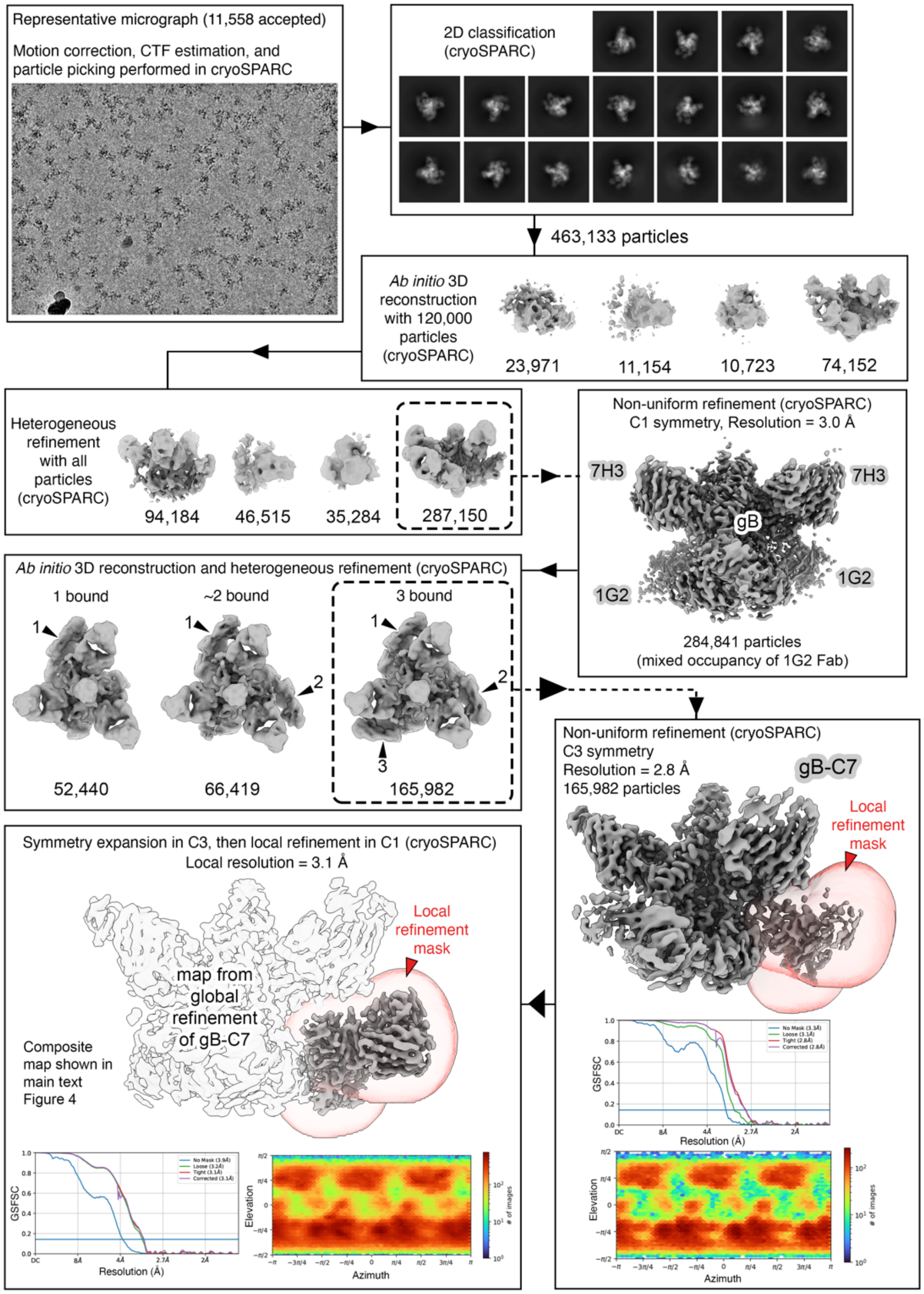


**Figure S6. Cryo-EM data collection and processing summary for the structure of gB-C7 in complex with 1G2 and 7H3 Fabs**. Cryo-EM processing workflow consisted of the following steps performed in cryoSPARC v4.2.0: motion correction, CTF estimation, particle picking, 2D classification, *ab initio* 3D reconstruction, iterative heterogenous refinement, homogenous refinement, and non-uniform refinement with C3 symmetry. A mask was created around domain I and 1G2 using ChimeraX and imported to cryoSPARC for local refinement with C1 symmetry using the C3 symmetry expanded particle stack. The number of particles used in each step is listed. The final, highest resolution reconstructions for the global and local refinements are shown at the bottom of the figure above gold-standard Fourier shell correlation and directional distribution plots.

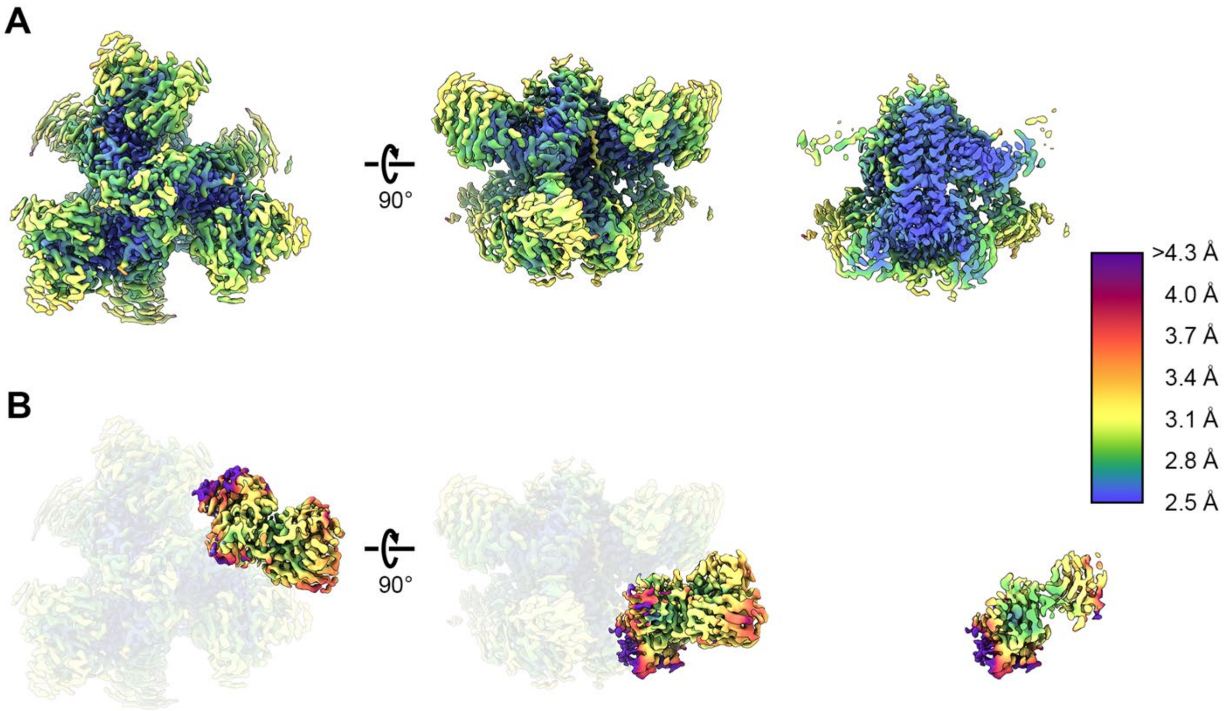


**Figure S7. Cryo-EM maps colored by local resolution from global and local refinement of gB-C7**. The (**A**) global and (**B**) local refinement maps of gB-C7 are colored as a rainbow according to the estimated local resolution from 2.5 Å (blue) to 4.3 Å (purple) resolution.

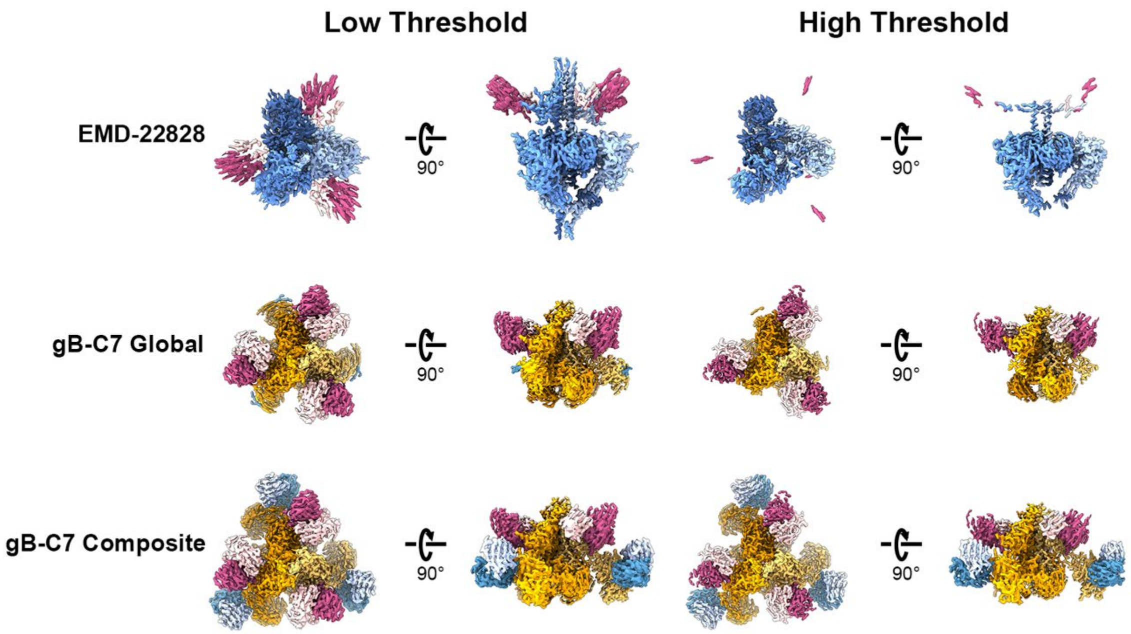


**Figure S8. Cryo-EM maps at different thresholds**. Top and side views of the previously determined EM map of prefusion HCMV gB (EMD-22828, PDB ID: 7KDP) (*23*) and the global and composite EM maps of HCMV gB-C7 complexed with 1G2 and 7H3 Fabs at lower and higher thresholds.

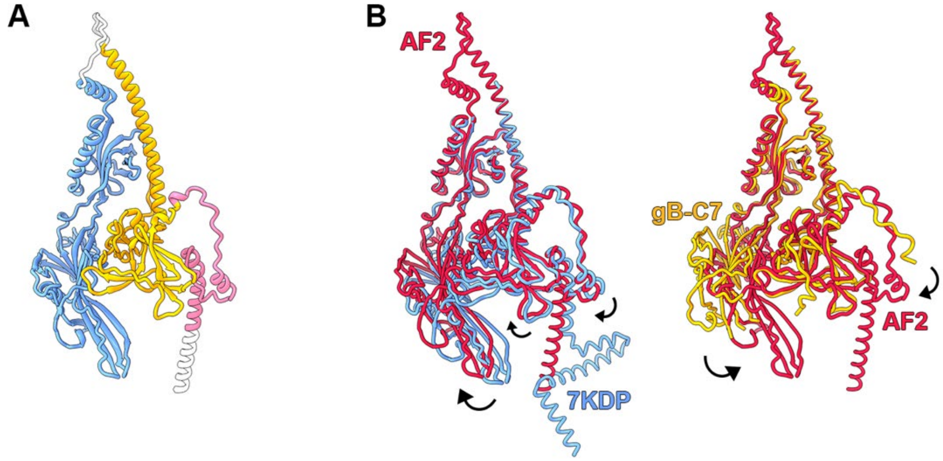


**Figure S9. AlphaFold2-predicted HCMV gB protomer in the prefusion conformation**. (**A**) AlphaFold2 (AF2) prefusion gB protomer model is shown as a ribbon diagram. The previously determined structure of prefusion HCMV gB (PDB ID: 7KDP) (*23*) was used as a template for AF2 prediction (*50*). The first and second regions expected to move during the conformational rearrangement from pre-to-postfusion gB are colored blue and pink, respectively, and the region that does not undergo substantial rearrangement is colored yellow. (**B**) The AF2 model (red) is superimposed with the previously determined structure of prefusion HCMV gB (blue, PDB ID: 7KDP) and the structure of gB-C7 (yellow), all shown as cartoon tube traces of the α-carbon backbones. Shifts in domain arrangement are highlighted with arrows.

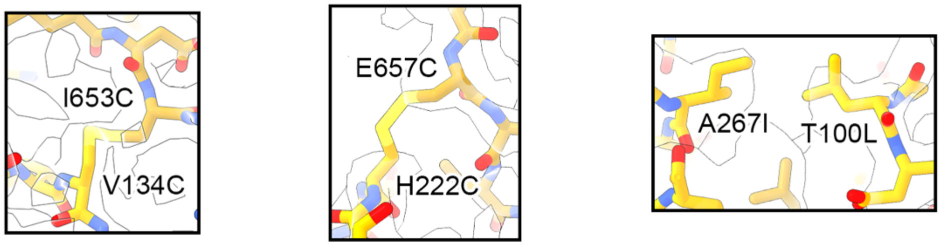


**Figure S10. gB-C7 substitutions supported by the cryo-EM map**. The model of HCMV gB combination variant gB-C7 is shown in gold sticks fit in the transparent composite EM map of the HCMV gB-C7 complex. Panels show close views of the substitutions that comprise design gB-C7: (**A**) V134C/I653C, (**B**) H222C/E657C, and (**C**) T100L/A267I. Sulfur atoms are shown in yellow, nitrogen atoms in blue, and oxygen atoms in red.

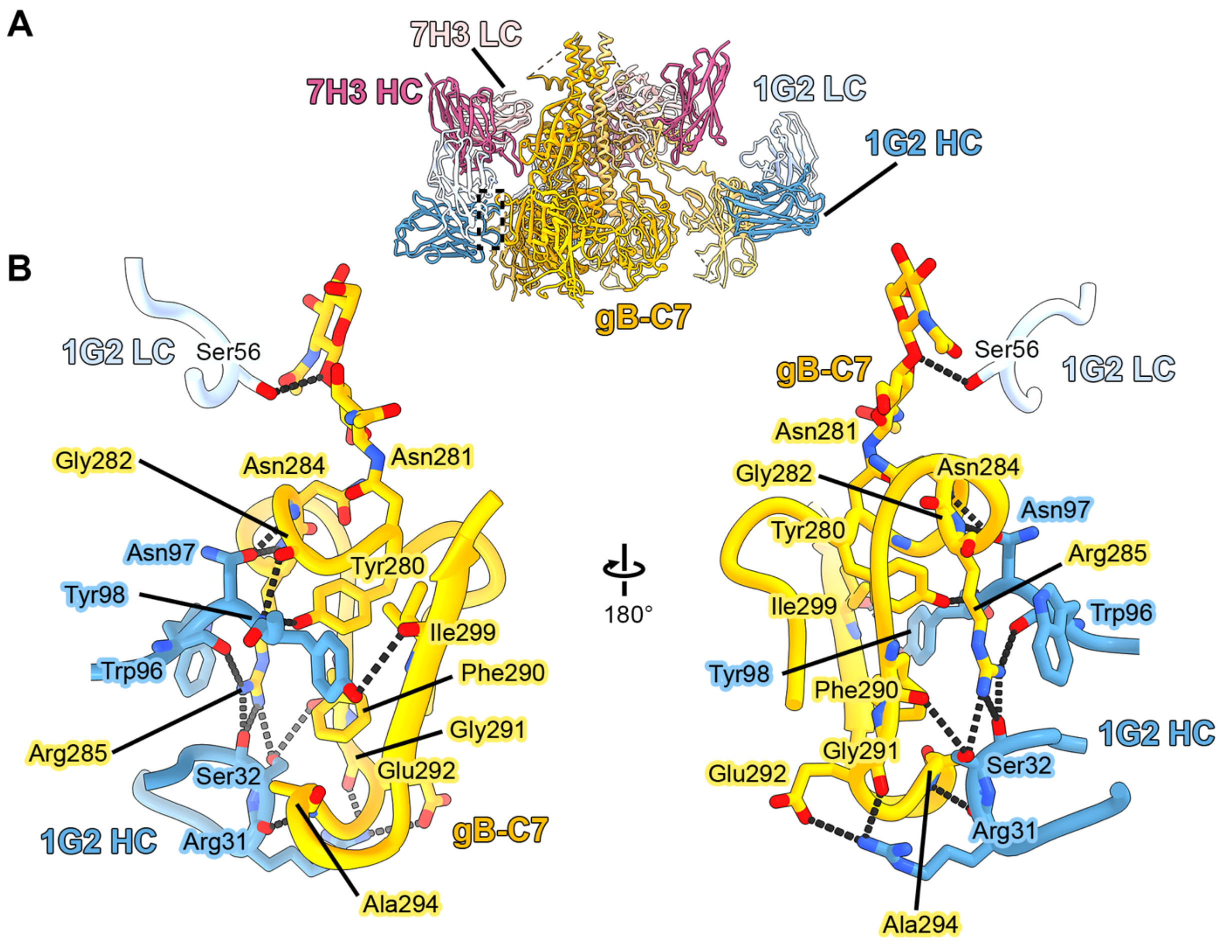


**Figure S11. The gB-C7:1G2 Fab interface**. (**A**) Model of gB-C7 (shades of yellow colored by protomer) complexed with 1G2 (heavy chain: blue, light chain: light blue) and 7H3 (heavy chain: pink, light chain: light pink) Fabs, with the region shown in B highlighted with dashed lines. (**B**) Zoomed view of the binding interface between 1G2 and gB-C7; the interface comprises 13 hydrogen bonds and 663 Å$^2$ of buried surface area on gB-C7. The gB-C7:1G2 interface is highly similar to the previously described postfusion gB:1G2 interface (PDB ID: 5C6T) (*24*) with an RMSD of 0.4 Å across the 21 Cα atoms comprising the continuous 1G2 epitope on gB-C7. Hydrogen bonds are shown as black dashes with interacting residues labeled and shown as sticks. Sulfur atoms are shown in yellow, nitrogen atoms in blue, and oxygen atoms in red.

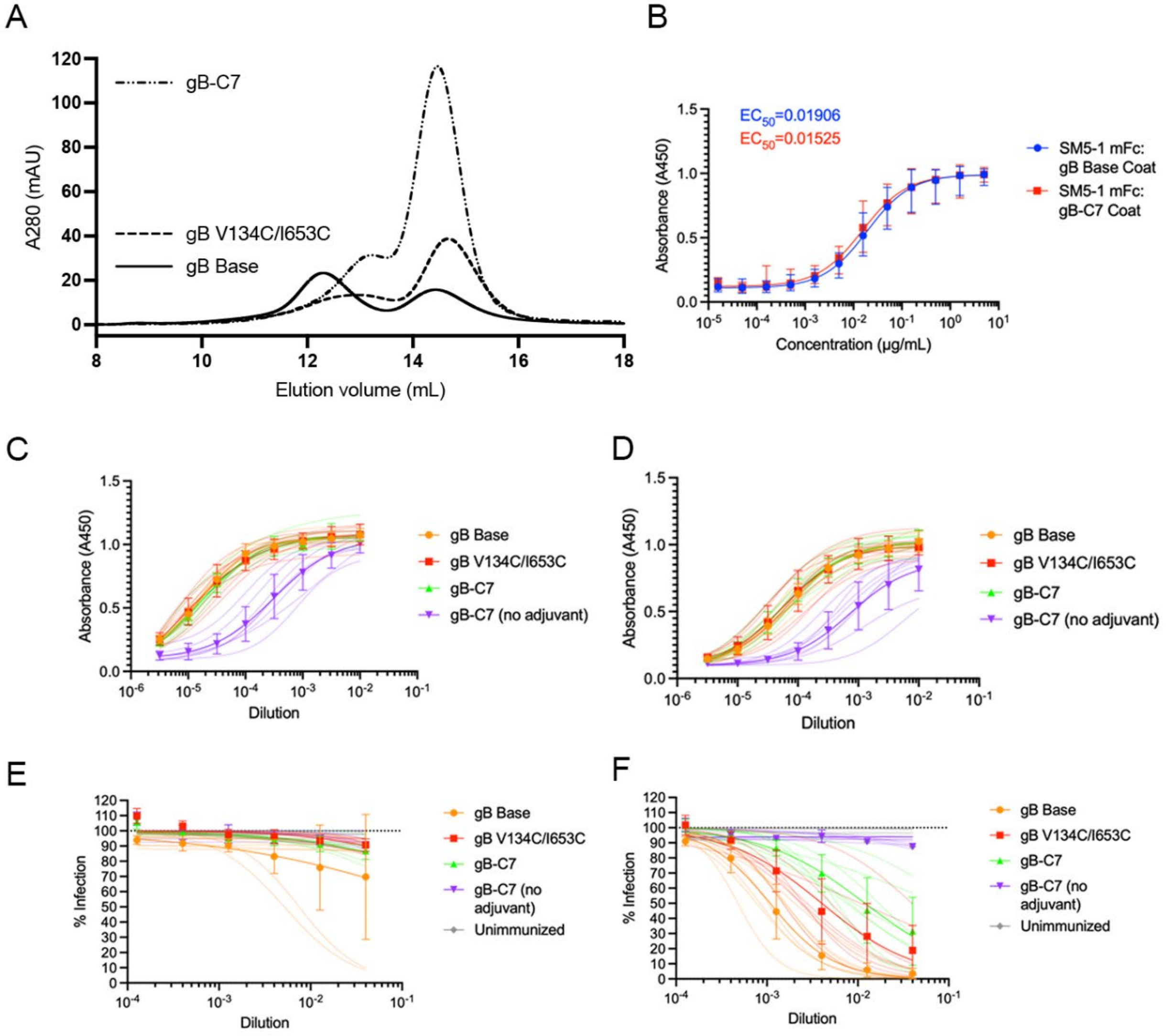


**Figure S12. Individual ELISA and neutralization curves for anti-gB mouse sera**. ELISA plates were coated with gB Base or gB-C7 and incubated with the SM5-1 murine IgG2a Fc (mFc) control or mouse serum. Bound mouse antibodies were detected using goat anti-mouse Ig HRP. (**A**) Size-exclusion chromatography (SEC) traces of purified gB constructs gB Base, gB V134C/I653C, and gB-C7 prepared for immunization. (**B**) SM5-1 mFc binds with similar affinity to gB Base and gB-C7 on ELISA. Plotted curves and error bars represent the average and standard deviation across 33 ELISA plates. Sera samples were diluted by √10 from 1 x $10^{-2}$–3.16 x $10^{-6}$ and incubated on ELISA plates coated with (**C**) gB Base or (**D**) gB-C7. Immunized mouse serum was incubated with AD169-GFP in a √10-fold dilution series from 4 x $10^{-2}$–1.3 x $10^{-4}$ (**E**) without or (**F**) with 12.5% guinea pig complement prior to incubation with human MRC-5 fibroblasts. Individual mice are represented by lighter-colored lines, which are averages across two independent experiments. The averages of each group are shown in heavier lines with error bars representing the standard deviation across each group ($n$ = 8 mice/group).

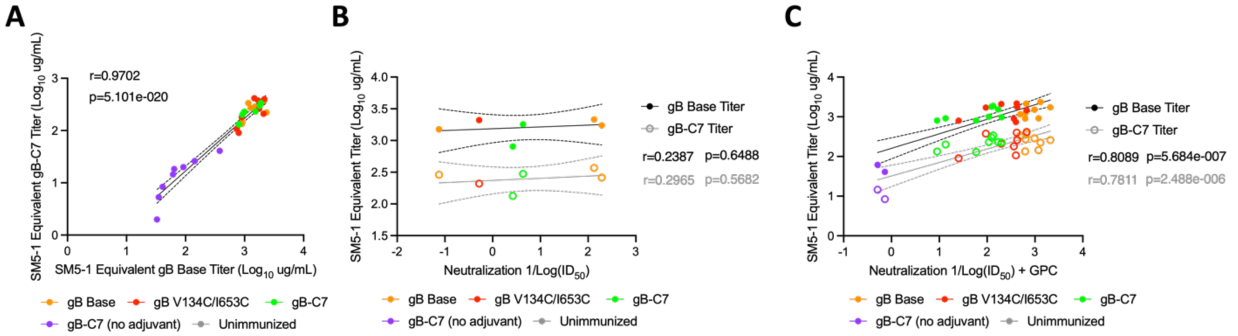


**Figure S13. Correlations between gB binding and neutralizing antibody titers**. Pearson's correlation coefficients (r) and p-values were calculated between (**A**) gB Base binding and gB-C7 binding antibody titers, (**B**) neutralization and gB Base binding or gB-C7 binding antibody titers, and (**C**) neutralization in the presence of 12.5% guinea pig complement and gB Base binding or gB-C7 binding antibody titers.

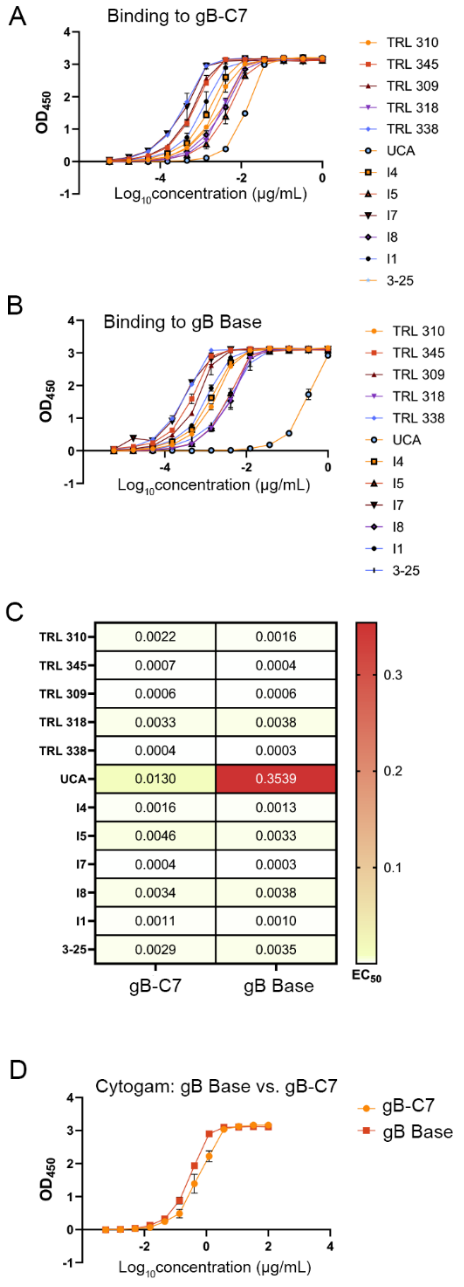


**Figure S14. The AD-2S1-specific TRL345 Unmutated Common Ancestor binds gB-C7 more potently than gB Base.** Binding of anti-gB AD-2 TRL345 lineage antibodies to (**A**) gB-C7 and (**B**) gB Base via ELISA. (**C**) ELISA plates were coated with gB Base or gB-C7 and incubated with the TRL345 lineage antibodies listed. Most of the TRL345 lineage antibodies exhibited similar binding to gB-C7 and gB Base. However, the TRL345 unmutated common ancestor (UCA) demonstrated higher binding to gB-C7 than to gB Base. (**D**) Cytogam was used as a positive control for binding gB-C7 and gB Base.

**Table S1.** Cryo-EM data collection and refinement statistics

| | | | |
|---|---|---|---|
| **CRYO-EM DATA COLLECTION** | | | |
| Microscope (FEI) | Titan Krios | Titan Krios | |
| Voltage (kV) | 300 | 300 | |
| Detector | K3 | K3 | |
| Pixel size (Å/pix) | 0.8332 | 0.8332 | |
| Exposure ($e^-/Å^2$) | 80 | 80 | |
| Defocus range (μm) | 1.0-3.0 | 1.0-2.5 | |
| Micrographs collected at 0° tilt | 1,308 | 12,524 | |
| Micrographs collected at 30° tilt | 7,371 | 0 | |
| Micrographs used (all tilt angles) | 7,939 | 11,558 | |
| Total Particles extracted | 4,317,580 | 3,590,995 | |
| Automation software | SerialEM 3.9.0 beta | SerialEM 3.9.0 beta | |
| **Sample Composition** | **postfusion gB Base in complex with 1G2 and 7H3** | **prefusion gB-C7 in complex with 1G2 and 7H3 Fabs** | |
| **CRYO-EM REFINEMENT STATISTICS** | | | |
| Particles | 211,132 | 165,982 | |
| Symmetry imposed | C1 | C3 | C1 |
| Map sharpening B-factor | 98 | 86 | 102 |
| Resolution at FSC... | | | |
| Unmasked: 0.5 (Å) | 6.0 | 3.8 | 4.6 |
| Masked: 0.5 (Å) | 3.8 | 3.2 | 3.7 |
| Unmasked: 0.143 (Å) | 4.0 | 3.3 | 3.9 |
| Masked: 0.143 (Å) | 3.4 | 2.8 | 3.1 |
| **MODEL REFINEMENT AND VALIDATION STATISTICS** | | | |
| Composition | | | |
| Amino Acids (#) | 2218 | 2958 | |
| Ligands (Type: #) | BMA: 3, NAG: 42 | BMA: 3, NAG: 30 | |
| RMSD Bonds | | | |
| Length [Å] (# > 4s) | 0.002 (0) | 0.004 (0) | |
| Angles [°] (# > 4s) | 0.44 (0) | 0.57 (0) | |
| Ramachandran plot | | | |
| Outliers (%) | 0.1 | 0.0 | |
| Allowed (%) | 3.8 | 5.0 | |
| Favored (%) | 96.1 | 95.0 | |
| Rotamer outliers (%) | 0.7 | 0.0 | |
| C-β outliers (%) | 0.0 | 0.0 | |
| CaBLAM outliers (%) | 2.0 | 1.6 | |
| CC (mask) | 0.87 | 0.87 | |
| MolProbity score | 1.6 | 1.5 | |
| Clash score | 6.3 | 3.3 | |
| Q-score | 0.48 | 0.55 | |
| PDB ID | 8VYM | 8VYN | |
| EMDB ID(s) | EMD-43667 | EMD- 43670, 43671, 43672 | |
| **Sample Composition** | **Postfusion gB Base in complex with 1G2 and 7H3** | **Prefusion gB-C7 in complex with 1G2 and 7H3 Fabs** | |

**Table S2.** Area under the curve from SEC traces for HCMV gB single substitution variants normalized to gB Base.

| Substitution(s) | Normalized $AUC_{LMW}$ | Normalized $AUC_{HMW}$ | Normalized $AUC_{LMW+HMW}$ | Normalized $AUC_{LMW/HMW}$ |
|---|---|---|---|---|
| Q98C/G271C | 0.14 | 0.46 | 0.20 | 3.35 |
| Q98C/N658C | 0.76 | 3.32 | 1.26 | 4.35 |
| V134C/I653C | 1.26 | 3.63 | 1.72 | 2.88 |
| N220C/E657C | 0.59 | 4.52 | 1.36 | 7.61 |
| H222C/E657C | 1.35 | 5.07 | 2.08 | 3.77 |
| I356C/A500C | 0.46 | 0.68 | 0.50 | 1.49 |
| S367C/A503C | 1.61 | 4.55 | 2.18 | 2.83 |
| Q612C/R662C | 0.57 | 1.42 | 0.74 | 2.47 |
| S674C/E698C | 1.82 | 2.98 | 2.04 | 1.64 |
| V273K | 1.61 | 1.66 | 1.62 | 1.03 |
| D509A | 0.38 | 0.51 | 0.40 | 1.36 |
| D509L | 0.05 | 0.11 | 0.06 | 2.30 |
| S526K | 0.94 | 1.30 | 1.01 | 1.38 |
| D679V/E682N | 1.84 | 2.29 | 1.96 | 1.24 |
| Q98L/N658I | 0.98 | 2.35 | 1.25 | 2.39 |
| T100L/A267I | 1.44 | 5.00 | 2.06 | 3.47 |
| K130Y | 1.75 | 4.02 | 2.33 | 2.29 |
| K260W | 1.11 | 2.53 | 1.36 | 2.27 |
| L664F | 1.54 | 1.92 | 1.61 | 1.25 |
| N341P | 2.03 | 1.53 | 1.94 | 0.75 |
| L484P | 1.72 | 4.09 | 2.14 | 2.38 |
| V645P | 0.95 | 2.03 | 1.17 | 2.14 |
| D646P | 0.98 | 1.81 | 1.12 | 1.85 |
| E686P | 1.78 | 1.25 | 1.69 | 0.70 |

AUC was quantified from SEC traces of purified gB single substitution variants. AUCs of the low molecular weight (LMW) peak ($AUC_{LMW}$), the high molecular weight (HMW) peak ($AUC_{HMW}$), and the sum of the two peaks ($AUC_{LMW+HMW}$) were normalized to respective gB Base AUCs expressed and purified within the same batch. The ratio of the normalized $AUC_{LMW}$ relative to the normalized $AUC_{HMW}$ ($AUC_{LMW/HMW}$) is reported at the far right.

**Table S3.** Anti-gB antibody titers and neutralization capacity of immunized mouse serum

| Immunogen | SM5-1 Equivalent Titer (µg/mL) | | Neutralization 1/$ID_{50}$ | |
|---|---|---|---|---|
| | gB Base Binding | gB-C7 Binding | No complement | With complement |
| gB Base | 1502.3 | 287.4 | 0.1 | 978.5 |
| | 1155.4 | 335.3 | ND | 492.4 |
| | 918.8 | 143.7 | ND | 1192.9 |
| | 1261.6 | 271.4 | ND | 635.7 |
| | 2152.3 | 370.3 | 133.4 | 659.6 |
| | 2352.2 | 222.6 | ND | 1300.2 |
| | 1729.8 | 259.8 | 193.7 | 2092.5 |
| | 902.0 | 134.0 | ND | 624.2 |
| gB V134C/I653C | 1703.4 | 374.8 | ND | 93.8 |
| | 896.8 | 179.2 | ND | 362.8 |
| | 1771.6 | 260.9 | ND | 425.5 |
| | 1461.3 | 414.9 | ND | 670.7 |
| | 2097.1 | 209.3 | 0.5 | 197.8 |
| | 738.3 | 106.6 | ND | 398.7 |
| | 2154.9 | 399.8 | ND | 415.3 |
| | 797.1 | 90.3 | ND | 25.4 |
| gB-C7 | 1000.5 | 229.4 | ND | 108.8 |
| | 805.6 | 129.8 | ND | 59.2 |
| | 1803.9 | 297.6 | 4.3 | 124.0 |
| | 976.0 | 224.3 | ND | 200.2 |
| | 1887.4 | 338.5 | ND | 133.5 |
| | 1533.0 | 231.6 | ND | 169.5 |
| | 804.4 | 133.6 | 2.7 | 9.0 |
| | 912.8 | 200.2 | ND | 13.5 |
| gB-C7 (no adjuvant) | 142.7 | 26.1 | ND | ND |
| | 35.1 | 5.3 | ND | ND |
| | 40.9 | 8.4 | ND | 0.7 |
| | 91.8 | 20.0 | ND | ND |
| | 32.9 | 2.0 | ND | ND |
| | 378.5 | 41.0 | ND | ND |
| | 64.2 | 18.3 | ND | ND |
| | 61.4 | 14.5 | ND | 0.5 |

Antibody binding titers against gB Base and gB-C7 of immunized mouse sera. Titers were measured by ELISA relative to SM5-1 mFc standard. The capacity of immunized mouse sera to neutralize AD169 infection of MRC-5 fibroblasts is reported as 1/$ID_{50}$. Data is reported as an average of two independent experiments. ND: Not detected.
